# Efficient Suppression of Endogenous CFTR Nonsense Mutations Using Anticodon Engineered Transfer RNAs

**DOI:** 10.1101/2021.10.09.463783

**Authors:** Wooree Ko, Joseph J. Porter, Matthew T. Sipple, Katherine M. Edwards, John D. Lueck

## Abstract

Nonsense mutations or premature termination codons (PTCs) comprise ∼11% of all genetic lesions, which result in over 7,000 distinct genetic diseases. Due to their outsized impact on human health, considerable effort has been made to find therapies for nonsense-associated diseases. Suppressor tRNAs have long been identified as a possible therapeutic for nonsense-associated diseases, however their ability to inhibit nonsense-mediated mRNA decay (NMD) and support significant protein translation from endogenous transcripts has not been determined in mammalian cells. Here we investigated the ability of anticodon edited (ACE)-tRNAs to suppress cystic fibrosis (CF) causing PTCs in the cystic fibrosis transmembrane regulator (*CFTR*) gene in gene-edited immortalized human bronchial epithelial (16HBEge) cells. Delivery of ACE-tRNAs to 16HBEge cells harboring three common CF mutations G542X-, R1162X- and W1282X-CFTR PTCs significantly inhibited NMD and rescued endogenous mRNA expression. Furthermore, delivery of our highly active leucine encoding ACE-tRNA resulted in rescue of W1282X-CFTR channel function to levels that significantly exceed the necessary CFTR channel function for therapeutic relevance. This study establishes the ACE-tRNA approach as a potential stand-alone therapeutic for nonsense-associated diseases due to its ability to rescue both mRNA and full-length protein expression from PTC containing endogenous genes.

**One Sentence Summary:** Suppression of endogenous CFTR nonsense mutations by anticodon engineered tRNAs significantly increases mRNA expression and channel function.

## INTRODUCTION

Nonsense mutations are single nucleotide mutations that convert a canonical amino acid codon to one of the three stop codons (UAA, UAG, UGA). Nonsense mutations result in a premature termination codon (PTC) generally with almost complete loss-of-function of the affected protein. This is due to both the PTC terminating translation prematurely, resulting in a truncated protein, and nonsense mediated mRNA decay (NMD) degrading the mRNA coding for the PTC-containing protein (1). With the advent of next-generation sequencing, more than 7,500 PTCs in nearly 1,000 different human genes have been discovered. PTCs account for close to 11% of all described mutations leading to inherited human disease (2). An abridged list of the known PTC-associated disease phenotypes includes Duchenne muscular dystrophy (3), spinal muscular atrophy (4), β-thalessemia (5), Hurler syndrome (6), Usher syndrome (7), and cystic fibrosis (8).

Cystic fibrosis (CF), a common autosomal recessive genetic disease, is caused by mutations to the cystic fibrosis transmembrane conductance regulator (*CFTR*) gene (9-11). Similar to genetic diseases as a whole, PTCs account for close to 10% of mutations causing CF (12). Several therapeutic small molecules rescue clinically relevant levels of CFTR function for a growing percentage of people with CF. However, all CF therapeutic small molecules so far target full-length aberrant CFTR protein (13-15). As PTCs result in little to no full-length protein, these small molecules targeting the CFTR protein are ineffective at rescuing PTC-containing CFTR. Instead, PTC therapies must target the defective gene, post-transcriptional processing, or processes governing protein translation.

Shortly after the discovery of the *CFTR* gene (9-11), proof-of-concept for gene replacement was demonstrated (16, 17). In the following 30 years, significant effort has been directed towards CFTR gene replacement therapy as this method has potential to repair the phenotypic defect for all people with CF. A number of viral and non-viral gene delivery approaches have been investigated (18). Improvements in vector design and delivery methods, a deeper understanding of lung biology and disease development, and new animal models have gotten us closer to the development of an effective mutation agnostic gene therapy for CF, however to date no gene replacement therapy has made it to widespread clinical use (18).

CRISPR/Cas-based technologies including prime editing, homologous recombination following double-strand breaks, and in-frame deletions have received considerable interest as potential nonsense mutation interventions (19-21). As with total gene replacement, the CRISPR/Cas approach requires efficient delivery, optimally to progenitor cells, to ensure long-term repair of the nonsense mutation. Delivery of CRISPR/Cas both as an AAV encoding the CRISPR/Cas pair (20) and as a CRISPR/Cas ribonucleoprotein delivered with engineered amphophilic peptides (21) has been demonstrated in model systems. A drawback to CRISPR/Cas approaches is that design of a new guide RNA is required for each mutation site. As with gene therapy, an efficient delivery method for clinical use has not been determined.

Small molecules including aminoglycosides, synthetic aminoglycosides, and oxodiazoles have been reported to act as nonsense readthrough agents (22-24). Pharmacological treatment resulting in PTC readthrough is an attractive therapeutic avenue to pursue as these promise to allow for body-wide readthrough of PTCs. Small molecule PTC readthrough agents generally bind to the ribosome and reduce translational fidelity. While this allows for suppression of PTCs by near-cognate tRNAs, global translational fidelity is also decreased. Of note, several of these small molecule PTC readthrough agents have been investigated in clinical trials with generally low efficacy and strong oto- and nephrotoxicity (25). In addition, a recent study has also indicated that genome-wide natural termination codon (NTC) readthrough stimulated by the aminoglycoside G418 *in vivo* results in the disruption of several biological processes with detrimental cellular effects (26). While the promises of small molecule nonsense readthrough agents are tantalizing, due to issues with safety, efficacy, or both, their clinical implementation has remained out of reach.

Another nonsense suppression technology that has been explored previously is the use of nonsense suppressor tRNAs or anticodon edited tRNAs (ACE-tRNAs) (5, 27-31). A recently developed library of ACE-tRNAs with the anticodons engineered via mutagenesis to suppress one of the UAA, UAG, or UGA PTC codons demonstrated efficacious *in vivo* PTC suppression (30). While previous work by ourselves and others has demonstrated the safe and efficacious ability of ACE-tRNAs to rescue PTCs in the context of cDNAs, it is still unknown if they can take a major step forward in their therapeutic potential and rescue PTCs in a native genomic context. Heterologous overexpression of PTC-containing mRNAs as cDNAs has fundamentally limited relevance, as these constructs lack the genomic context/chromatin environment, native protein promoter, intronic sequences and post-transcriptional processing.

To that end, we set out to demonstrate ACE-tRNA based rescue of PTC-containing CFTR from the well characterized immortalized isogenic human bronchial epithelial (16HBE14o-) cell lines modified with CRISPR/Cas9 to contain the most common CF-causing PTC variants (32). Here we show significant rescue of PTC-containing CFTR mRNA transcripts by real-time qRT-PCR for three of the most common CF-causing PTCs (G542X (2.5%), W1282X (1.2%), and R1162X (0.4%)) (12) and significant rescue of CFTR protein function by patch clamp electrophysiology following delivery of our best performing ACE-tRNA^Leu^. The level of ACE-tRNA dependent rescue of functional W1282X-CFTR protein demonstrated here, is well above the predicted mark for therapeutically meaningful rescue (33, 34). While efficient delivery of ACE-tRNAs to affected cell types remains a hurdle to their therapeutic implementation, ACE-tRNAs display several attractive qualities as gene therapy agents bolstering their therapeutic promise (31). The results presented here display an exciting step forward in an unmet medical need for patients with nonsense mutations.

## RESULTS

### Development of a nonsense suppression reporter system

While nonsense mutations cause several serious genetic diseases, here we chose to focus on CF, using the 16HBE14o-cell line and the derivative 16HBE gene edited (16HBEge) cell lines with engineered R1162X-, W1282X-, or G542X-CFTR variants (32). The genome of each of these 16HBEge cell lines was modified via CRISPR/Cas9 to contain a TGA stop codon at the amino acid residue specified in the *CFTR* gene. This model retains the native CFTR transcriptional, post-transcriptional, and translational regulation. We first set out to understand which previously identified nonsense suppressor ACE-tRNAs, whether delivered as DNA or RNA, were most effective. We also wanted to determine their persistence of activity and if previous results were recapitulated in human epithelial cells (30). To do this, we wanted a flexible high-throughput method to assess nonsense suppression in different cell lines, which could be transferred to any cell line of interest for future studies. Here we chose to use the *piggyBac* (PB) transposon/transposase system, which integrates the gene of interest (or cargo) specifically into genomic TTAA sites, facilitates fast selection of stable cell lines, can be used to generate both cell lines and animal models, and for which a hyperactive transposase has been generated (Fig. 1A) (35). We used a PTC-containing NanoLuc (NLuc) luciferase as a nonsense suppression reporter due to high signal-to-noise and utility in high-throughput assays(30). We sought an assay to determine the potentially short half-lives of ACE-tRNA therapeutic agents (i.e. delivered as RNA) for this we appended the protein-destabilizing PEST domain (Fig. 1B), which has been previously shown to shorten the half-life of NLuc to ∼20 minutes in several cell lines (36). Traditional NLuc assays require cell disruption as NLuc is retained intracellularly (36). By appending the IL6 secretion signal to the NLuc reporter (Fig. 1B), we developed a method to assay nonsense suppression longitudinally without disrupting cells. For longitudinal nonsense suppression studies the cell media is assayed for PTC-containing Sec-NLuc luminescence (Fig. 1E). Further Our best performing ACE-tRNA^Gln^, ACE-tRNA^Glu^, ACE-tRNA^Arg^, ACE-tRNA^Gly^, ACE-tRNA^Leu^, ACE-tRNA^Trp^ library members displayed robust nonsense suppression for all three NLuc formats in 16HBE14o-cells, both as a transient co-transfection (Fig. 1C-E, Fig. S1A-C) and following reporter stable integration (Fig. 1F, Fig. S1D). These results closely track what was displayed previously in HEK293T cells, indicating there is no overt cell-type-dependent impact on ACE-tRNA activity.

**Figure 1.**
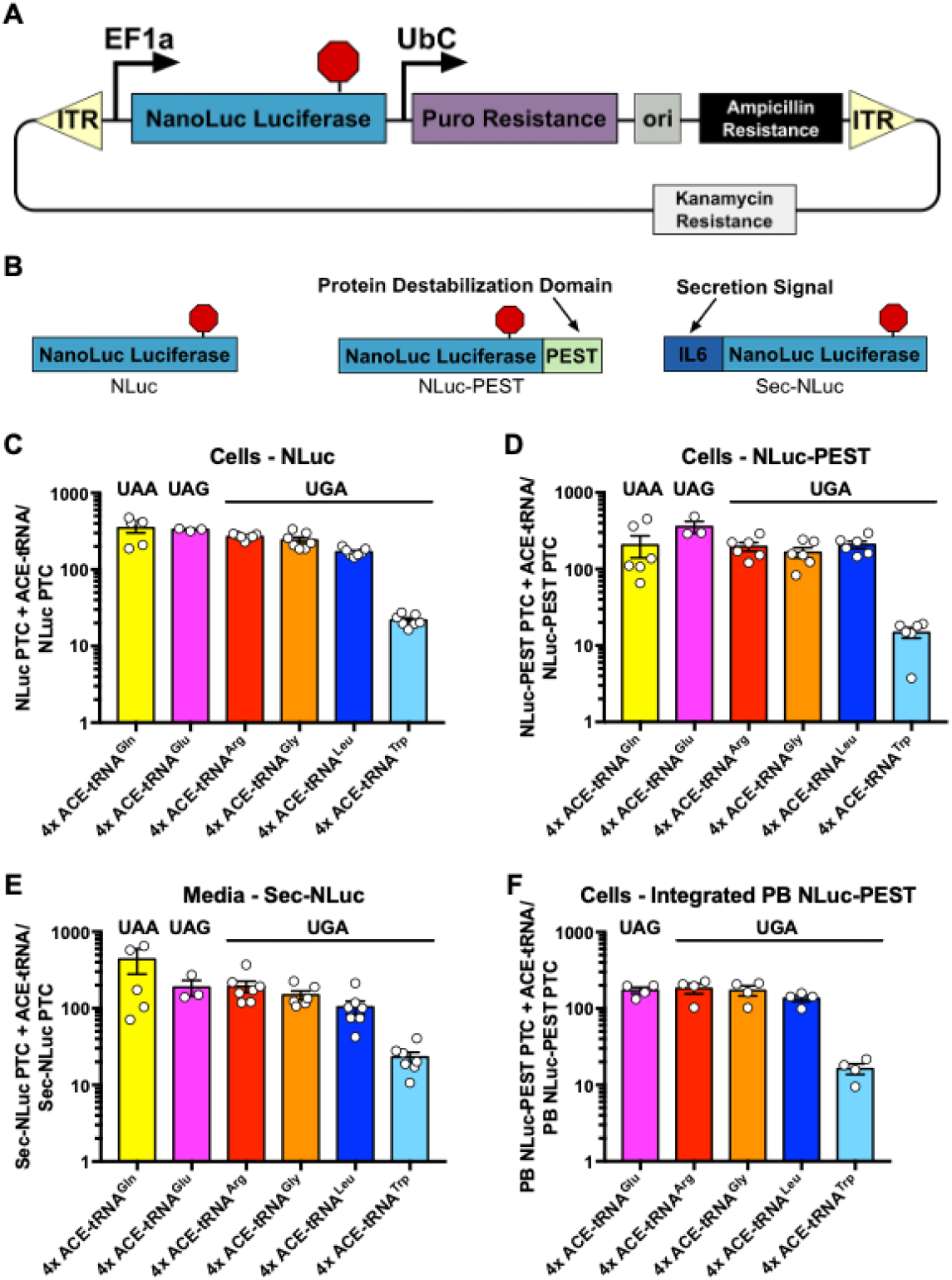
NLuc PTC reporter platforms faithfully report PTC suppression efficiency of ACE-tRNAs in 16HBE14o-cells. (**A**) Schematic illustrates the *piggyBac* (PB) system used in this study. (**B**) Three types of NLuc PTC reporters were generated for reporting either UAA, UAG, or UGA PTC suppression efficiency of ACE-tRNAs: NLuc, NLuc with protein destabilization domain (NLuc-PEST) and NLuc with secretion signal (Sec-NLuc). The wildtype NLuc PTC reporter allows for high-throughput screening of different ACE-tRNAs, the NLuc-PEST PTC reporter with short intracellular lifetime allows for the measurement of the half-life of ACE-tRNA and the Sec-NLuc PTC reporter with secretion capacity allows for longitudinal studies. (**C-F**) In WT 16HBE14o-cells co-transfected with ACE-tRNAs and PTC reporters, six highest performing ACE-tRNAs suppressing UAA, UAG and UGA PTCs exhibit a similar trend of luminescence rescue for (**C**) NLuc, (**D**) NLuc-PEST and (**E**) Sec-NLuc reporters. (**F**) The NLuc-PEST PTC reporter stably integrated in WT 16HBE14o-cells using the PB system are capable of reporting PTC suppression efficiency of transiently transfected ACE-tRNAs through luminescence. Data are presented as normalized mean luminescence ± SEM. *n* = 3-7, with each replicate represented as a circle. Raw luminescence of described experiments is presented in **Supplementary Fig. 1**.

### PTC suppression by ACE-tRNA encoded in plasmid peaks 48 hours following delivery to 16HBEge cells

To assess the kinetics of nonsense suppression rescue, we used R1162X- and W1282X-CFTR 16HBEge cells stably integrated with PB transposon containing Sec-NLuc UGA. The cells were seeded on Transwells and transfected with a plasmid containing 4 copies of our top performing ACE-tRNA^Arg^, ACE-tRNA^Trp^, and ACE-tRNA^Leu^ UGA suppressors, or an empty plasmid (control). Following transfection, the cells were seeded on Transwells and the Sec-NLuc luminescence was assayed daily from media collected from both the apical and basolateral sides of the Transwell. ACE-tRNA^Arg^ supported robust rescue of Sec-NLuc luminescence peaking around 2 days after transfection, in contrast to ACE-tRNA^Trp^, which performed relatively poorly (Fig. 2A-D). Indeed, the result with ACE-tRNA^Trp^ is not altogether unexpected, as it was the poorest performing ACE-tRNA in all of the most active ACE-tRNAs identified in our screen (*30*). The W1282L mutant of CFTR has been previously shown to support near-WT channel function (37), prompting us to test our highly performing ACE-tRNA^Leu^ for rescue of Sec-NLuc UGA in this assay. As with ACE-tRNA^Arg^, ACE-tRNA^Leu^ supported robust rescue of Sec-NLuc luminescence peaking 2 days after transfection (Fig. 2A-D). Based on the results of this longitudinal study, it is likely that the decrease in luminescence rescue is based both on silencing and dilution of plasmids due to cell splitting (38, 39). Importantly, the luminescence-based assay informed us on when to perform more in-depth studies on CFTR mRNA expression and channel function described in this study.

**Figure 2.**
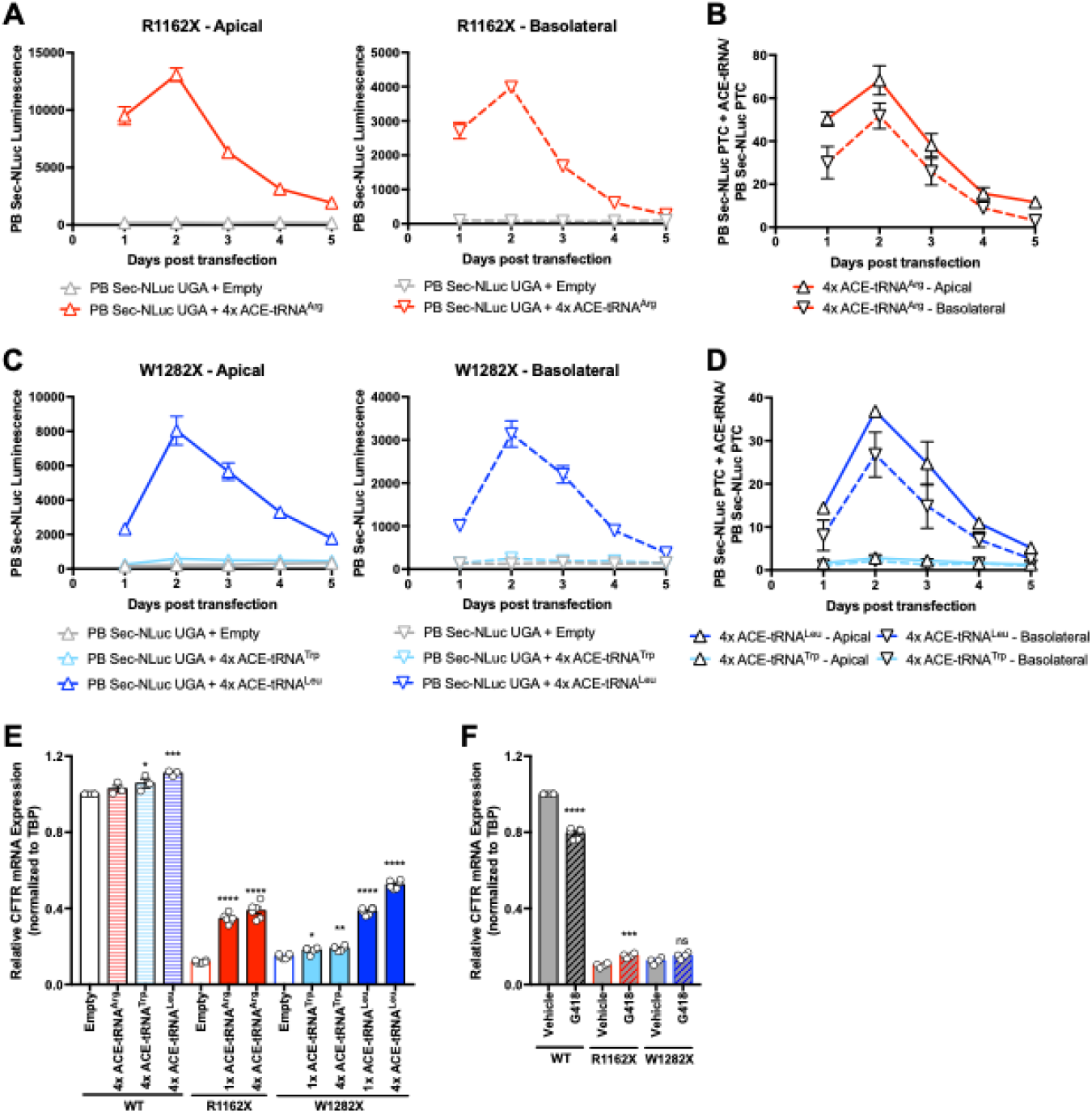
ACE-tRNAs delivered as plasmids have transient PTC suppression activity that significantly and robustly rescues endogenous R1162X- and W1282X-CFTR mRNA expression. (**A-B**) R1162X-CFTR and W1282X-CFTR cells stably expressing Sec-NLuc UGA reporter (R1162X PB Sec-NLuc UGA and W1282X PB Sec-NLuc UGA cells, respectively) were transfected with empty or 4x ACE-tRNA plasmids and luminescence measurements were made every 24 hrs from of media 5 consecutive days. Transfection of (**A, B**) ACE-tRNA^Arg^ (solid red, *n* = 3), (**C, D**) ACE-tRNA^Leu^ (dark blue, *n* = 3) and (**C, D**) ACE-tRNA^Trp^ (light blue, *n* = 3) plasmids results in time-dependent PTC suppression, with the peak luminescence detected at 2 days post transfection, in both the apical (solid lines) and basolateral (dashed lines) sides of Transwells. (**A-C**) Transfection of empty vectors (grey lines, *n* = 3) results in negligible luminescence change over the course of 5 days. Data are shown in raw luminescence. (**A, C**) Raw luminesce and (**B, D**) luminescence normalized to cells receiving empty plasmids (ACE-tRNA/Empty) display similar time-course of ACE-tRNA PTC suppression activity. (**E**) CFTR mRNA abundance normalized to TBP (housekeeping gene) in WT (white, *n* = 3), R1162X-CFTR (red, *n* = 6) and W1282X-CFTR (blue, *n* = 6) cells 48hrs following transfection with empty (open) or 1x and 4x ACE-tRNA (filled) plasmids (**E**) Suppression of R1162X- and W1282X-CFTR, with ACE-tRNA^Arg^ and ACE-tRNA^Leu^ or ACE-tRNA^Trp^, respectively, significantly recovers the level of CFTR transcripts 48hrs after delivery as determined by real-time qRT-PCR. (**F**) Treatment with G418 (100 μM), significantly increases CFTR mRNA expression in R1162X-CFTR cells (red) but not in W1282X-CFTR cells (blue). G418 treatment (gray, diagonal lines) significantly decreases CFTR mRNA expression in WT cells compared to vehicle treated cells (gray, open). Data are presented as average ± SEM. Significance was determined by (**E**) one-way ANOVA and Tukey’s post-hoc test and (**F**) unpaired t-test, where \**P*<0.05, \*\**P*<0.01, \*\*\**P*<0.001, and \*\*\*\**P*<0.0001.

### ACE-tRNAs rescue mRNA expression from endogenously encoded nonsense mutations

The CFTR transcripts in R1162X- and W1282X-CFTR 16HBEge cells are sensitive to degradation through the NMD pathway (1). To that end, we determined the level of NMD inhibition elicited by transfection of ACE-tRNA plasmids. Total RNA was isolated from R1162X- and W1282X-CFTR 16HBEge cells 2 days after transfection of either 1-copy or 4-copy ACE-tRNA^Arg^, ACE-tRNA^Trp^, ACE-tRNA^Leu^ plasmids. We then performed real-time qRT-PCR with probes against CFTR and control TATA-binding protein (TBP). ACE-tRNA^Arg^ and ACE-tRNA^Leu^, when delivered as plasmids containing 1-copy, rescued significant amounts of CFTR mRNA (34.6 ± 1.0 and 38.5 ± 0.8% of WT, respectively) as compared to cells transfected with an empty plasmid (11.9 ± 0.3 and 14.8 ± 0.4% of WT, respectively; Fig. 2E). Cells transfected with 4-copy ACE-tRNA plasmids also rescued significant amounts of CFTR mRNA (38.8 ± 1.7 and 52.5 ± 0.9% of WT, respectively), and to a greater extent than single copy ACE-tRNA plasmids, suggesting multiple ACE-tRNAs within each delivered vector is beneficial in non-saturating conditions (Fig. 2E), which is consistent with previously reported results (40).

ACE-tRNA dependent increase in CFTR mRNA expression is likely due to inhibition of NMD-dependent mRNA degradation by suppressing the PTC during pioneer rounds of mRNA translation (1). In contrast to ACE-tRNA treatment, G418 (100uM) treatment significantly decreased WT CFTR mRNA abundance in 16HBE14o-cells (79.5 ± 1.1% of vehicle treated 16HBE14o-cells), which is similar to a previously reported study (32). Reduction of WT CFTR mRNA expression following G418 treatment is likely due to global readthrough of NTCs and general translation miscoding (26), a feature that is not exhibited by ACE-tRNAs. Furthermore, treatment with G418 exhibited only a meager rescue of R1162X-CFTR mRNA expression (19.1 ± 0.5% of WT) compared to vehicle control (12.5 ± 0.8% of WT, Fig. 2F). To our knowledge, this is the first time that delivery of nonsense suppressor ACE-tRNAs has been directly shown to inhibit NMD and rescue mRNA from nonsense mutations in their endogenous genomic context.

### ACE-tRNA^Arg^ delivered as RNA suppresses nonsense mutations and rescues CFTR mRNA

We next moved to ascertain whether ACE-tRNAs delivered as RNA could also suppress nonsense mutations in 16HBEge cells. As with plasmid delivery, we first wanted to determine the time-course of rescue in cells for ACE-tRNA as RNA delivered by nucleofection. Use of the stable PB Sec-NLuc UGA reporter 16HBEge cell line displayed significant rescue at day 1 but decreased sharply at day 2 (Fig. S2A-D). As we missed the initial increase in Sec-NLuc luminescence observed previously with cDNA delivery, we likely under-sampled and missed the peak PTC suppression time-point for RNA delivery. With that in mind, we shifted to the NLuc-PEST assay as NLuc with an appended PEST sequence has a half-life of about 20 minutes (36). The activity of ACE-tRNA^Arg^ as RNA peaks at ∼5 hours and declines over the next 25 hours with a half-life of ∼6.5 hours (Fig. 3A, Fig. S2E). Using this time-course as a guide, we assayed the ability of ACE-tRNA^Arg^ delivered as RNA to inhibit NMD at both 6 and 24 hours after nucleofection. We found R1162X-CFTR mRNA expression was significantly rescued by ACE-tRNA^Arg^ RNA (31.9 ± 3.7% of WT) compared to mock (no RNA) nucleofections (13.0 ± 0.6% of WT) at 6 hours post nucleofection, however this effect was not observed at 24 hours post nucleofection (Fig. 3B). ACE-tRNA delivered as RNA convincingly suppresses endogenous nonsense mutations, however to become a viable therapy we will likely need to extend the half-life of ACE-tRNA activity or perform frequent delivery. It also may be the case that the relatively short ACE-tRNA RNA half-life may be a product of the target cells, where an extended ACE-tRNA RNA may be observed in cells that are not actively undergoing mitosis (i.e airway epithelial cells *in vivo*). Transient delivery of ACE-tRNAs as both cDNA and RNA resulted in robust NMD inhibition at early time-points, unfortunately the time-course of PTC suppression activity did not allow us to measure ACE-tRNA dependent rescue CFTR channel activity using the Ussing chamber measurements because on average a minimum of 4 days is needed for formation of tight junctions (32). We therefore pursued the possibility of generating 16HBEge cells with stably integrated ACE-tRNA expression cassettes for persistent PTC activity.

**Figure 3.**
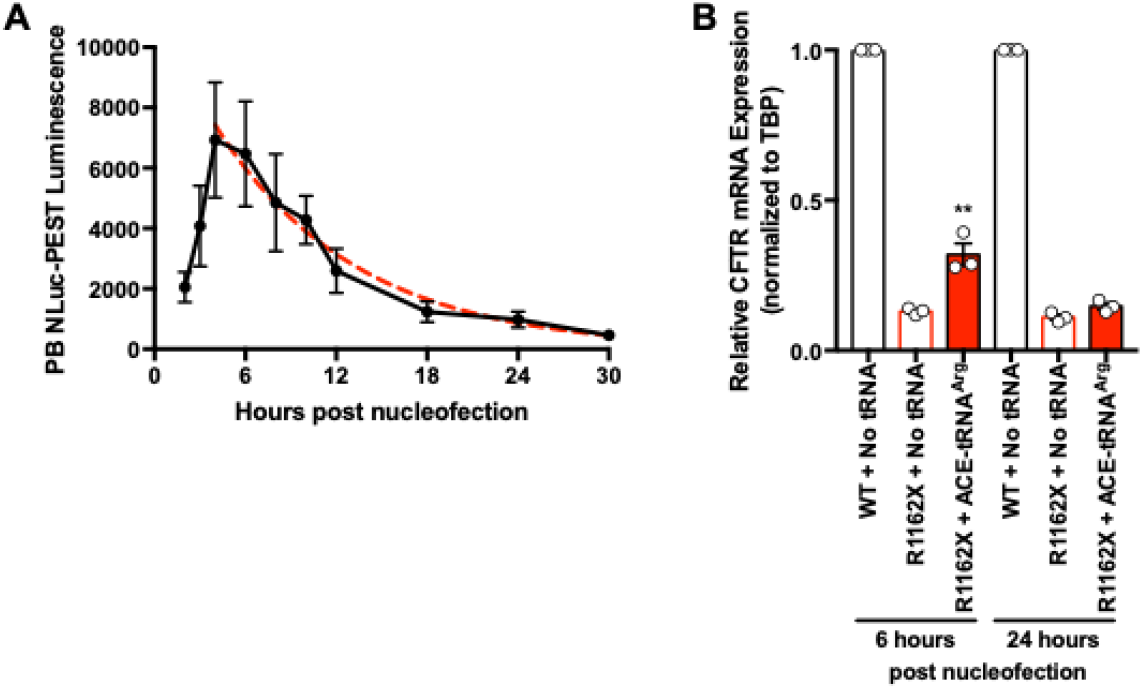
Delivery of ACE-tRNA^Arg^ as RNA to R1162X-CFTR 16HBEge cells results in significant rescue of CFTR mRNA expression. (**A**) R1162X-CFTR cells stably expressing NLuc-PEST UGA reporter (R1162X PB PEST-NLuc UGA cells) were nucleofected with and without ACE-tRNA^Arg^ RNA. Nonsense suppression mediated by ACE-tRNA^Arg^ was detected by luminescence measurements at 2, 3, 4, 6, 8, 10, 12, 18, 24 and 30 hours post nucleofection. Based on the average decay phase of luminescence by subtracting luminescence measurement with no delivered tRNA from that with ACE-tRNA^Arg^ RNA (red dotted line, y = 11413 e^-0.11x^), half-life of ACE-tRNA^Arg^ is 6.5 ± 0.3 hours. Raw luminescence measurements are presented in **Supplementary Fig. 2E**. (**B**) After 6 hours (left) of delivery, ACE-tRNA^Arg^ (red filled) delivered as RNA in R1162X-CFTR cells is sufficient to significantly rescue R1162X-CFTR mRNA expression as determined by real-time qRT-PCR, from no RNA control (red open). No significant rescue of R1162X-CFTR mRNA expression was observed at 24hrs (right). Data are presented as average ± SEM. All experiments contain an *n* = 3. Significance was determined by unpaired t-test, where \*\**P*<0.01.

### Stable genomic integration of ACE-tRNA^Arg^ rescues significant amounts of CFTR mRNA and channel function in R1162X-CFTR 16HBEge cells

Use of NLuc reporters allowed us to determine with ease a short time of action in both cDNA and RNA, likely due to rapid cell splitting of this model and dilution of ACE-tRNA plasmid. To circumvent this issue, we generated R1162X-CFTR 16HBEge cells containing stably integrated ACE-tRNA^Arg^ expression cassettes. For this purpose, we cloned 16 copies of ACE-tRNA^Arg^ along with a puromycin resistance marker into a PB transposon (Fig. S3A). R1162X-CFTR 16HBEge cells stably expressing PB ACE-tRNA^Arg^ were generated in a similar manner to the stable PB NLuc nonsense reporters described in Fig. 1 (see materials and methods). Of note, all studies with R1162X-CFTR 16HBEge cells stably integrated with 16 copies of ACE-tRNA^Arg^ transposon (R1162X PB 16x ACE-tRNA^Arg^ cells) were done from the total population of puromycin resistant cells which started from >100 colonies. We chose this approach to eliminate issues with genome insertion position effects of isogenic cell lines, which could pose issues with deconvoluting cell-to-cell variability. Given our ability to select 16HBEge cells stably expressing ACE-tRNA^Arg^, it assuages concerns that persistent expression of nonsense suppressor tRNAs at therapeutic levels will result in toxicity. Indeed, we observed that cells selected with puromycin exhibited similar morphology and cell division rates over >20 passages as compared to 16HBEge and 16HBE14o-cells, demonstrating that persistent ACE-tRNA expression is not toxic. To confirm that the stably incorporated ACE-tRNAs were functional, we transfected R1162X PB 16x ACE-tRNA^Arg^ cells with an NLuc UGA reporter construct. As expected, R1162X PB 16x ACE-tRNA^Arg^ cells exhibited significant (∼600 fold) NLuc PTC suppression (Fig. 4A, filled red bar) as compared to stable R1162X-CFTR 16HBEge cells generated with a transposon containing puromycin resistance without ACE-tRNAs (R1162X PB Empty; Fig. 4A, open red bar).

**Figure 4.**
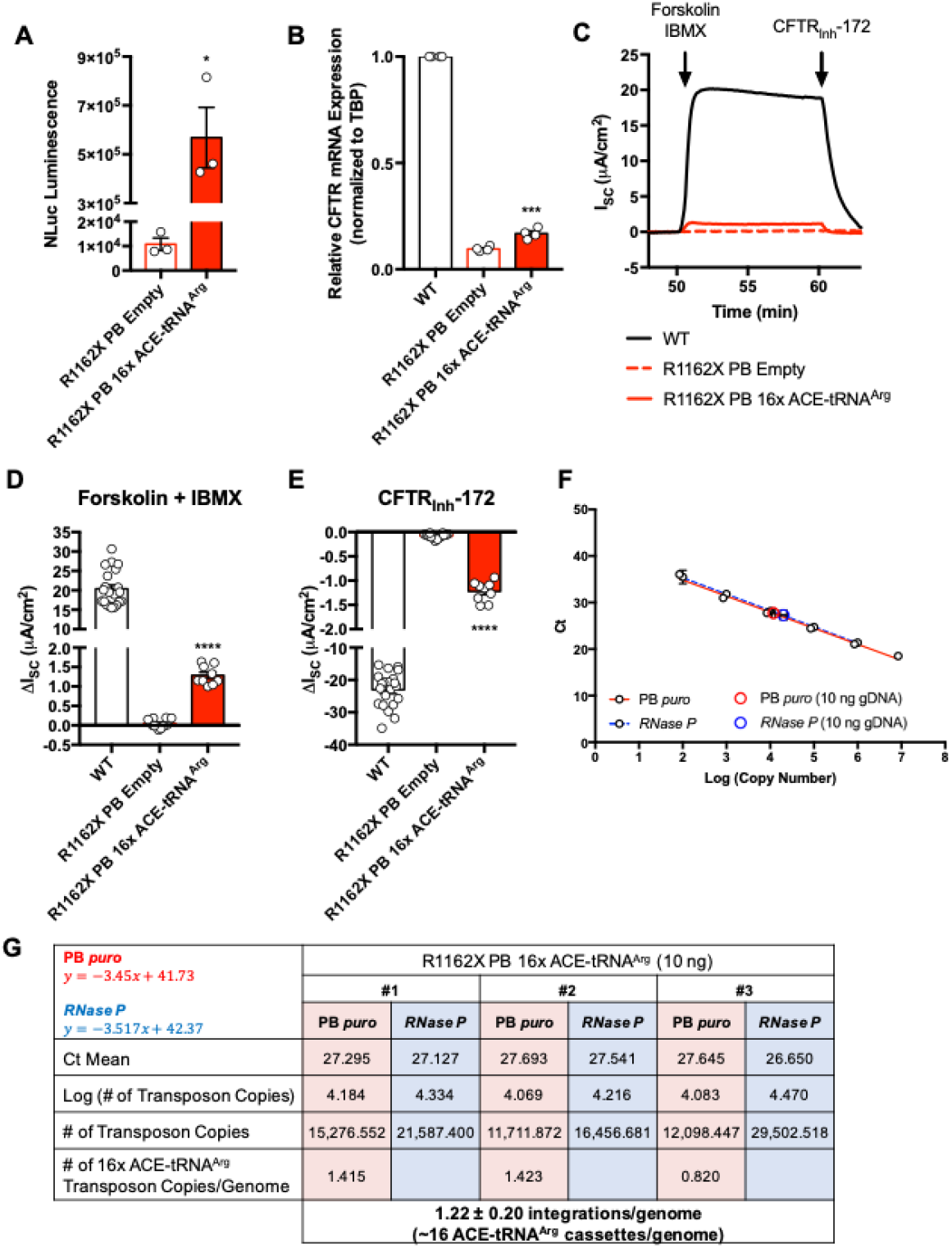
Stable integration of ACE-tRNA^Arg^ expression cassettes in R1162X-CFTR cells significantly rescues endogenous CFTR mRNA expression and CFTR channel function. (**A**) Raw luminescence measurement following transfection of the NLuc UGA reporter plasmid in R1162X-CFTR cells integrated with empty transposon (R1162X PB Empty; open red bar, *n* = 3) and R1162X-CFTR cells stably integrated with 16x ACE-tRNA^Arg^ transposon (R1162X PB 16x ACE-tRNA^Arg^; filled red bar, *n* = 3) results in significant suppression activity of integrated ACE-tRNA^Arg^. (**B**) R1162X-CFTR mRNA expression in R1162X PB 16x ACE-tRNA^Arg^ cells (filled red bar, *n* = 4)) is significantly increased compared to R1162X PB Empty cells (open red bars, *n* = 4). (**C**) Representative short-circuit Cl^-^ current traces recorded from WT (black trace), R1162X PB Empty (dashed red trace) and R1162X PB 16x ACE-tRNA^Arg^ (solid red trace) cells in response to forskolin (10 μM) and IBMX (100 μM), followed by CFTR inhibitor, CFTRInh-172 (20 μM). Average maximum ISC change in response to addition of (**D**) forskolin and IBMX and (**E**) CFTRInh-172 of WT 16HBE14o- (white bar, *n* = 21), R1162X PB Empty (open red bar, *n* = 14) and R1162X PB 16x ACE-tRNA^Arg^ (solid red bar, *n* = 9) cells. (**F**) qPCR standard curves generated from 10-fold dilutions of PB 16x ACE-tRNA^Arg^ (red line) and RNase P (Hs_RPP30 Positive Control; dashed blue) cDNA. qPCR measurements of PB *puro* transgene (open red circle) and endogenous *RNase P* gene (open blue circle) from 10ng input of gDNA from R1162X PB 16x ACE-tRNA^Arg^ cells. Each sample was assayed in triplicate by qPCR. (**G**)16x ACE-tRNA^Arg^ transgene copy number per genome in R1162X PB 16x ACE-tRNA^Arg^ cells is calculated by plotting Ct value to the standard curves. Data are presented as average ± SEM. Significance was determined by unpaired t-test, where \**P*<0.05, \*\*\**P*<0.001, and \*\*\*\**P*<0.0001.

We performed real-time qRT-PCR of total RNA isolated from R1162X PB 16x ACE-tRNA^Arg^ cells noting a significant albeit modest ∼7% increase (16.8 ± 1.2% of WT) in CFTR mRNA as compared to mRNA from R1162X PB Empty cells (9.6 ± 0.5% of WT; Fig. 4B). In the face of this modest rescue of mRNA, we determined the extent of R1162X-CFTR functional channel rescue. To do so, we used Ussing chamber recordings of CFTR chloride short circuit currents (32). Representative traces display a forskolin and 3-isobutyl-1-methlyxanthine (IBMX) response over baseline and that this response is blocked by CFTR_Inh_-172 (Fig. 4C), with average forskolin/IBMX and CFTR_Inh_-172 responses of CFTR channel function from R1162X PB 16x ACE-tRNA^Arg^ cells ranging from 5-7% of WT CFTR channel function from 16HBE14o-cells (Fig. 4D). Using the R1162X PB 16x ACE-tRNA^Arg^ cells, we can now determine for the first time how many ACE-tRNA^Arg^ expression cassettes are needed to support significant rescue of CFTR mRNA expression and channel function. Previous studies demonstrated on average 1-3 PB transposon integrations per genome (41, 42). To count the average number of ACE-tRNA^Arg^ transposon integration events per cell, we used qPCR to compare signals from the integrated transposon to standard curves generated from known quantities of target cDNA. The *RPP30* gene, subunit P30 of RNaseP enzyme complex, was used as a genomic DNA (gDNA) internal copy control, as this is a widely-accepted one copy gene of the haploid human genome (43). gDNA from R1162X PB 16x ACE-tRNA^Arg^ cells was isolated, and 10 ng was used as a template for real-time qPCR. The cycle threshold (C_t_) values obtained from primers and probes specific for PB *puro* (PB transposon puromycin resistance marker) and the *RPP30* gene (*RNase P*) revealed that on average only ∼1 copy of the PB transposon was integrated per genome, and therefore only 16 ACE-tRNA^Arg^ expression cassettes are integrated per genome (Fig. 4F, G).

Further experiments with 1 ng of gDNA or a probe targeted against the PB backbone using either 10 ng or 1 ng of gDNA revealed a similar average number of integration events (Fig. S3B-D). Indeed, we were excited to find that relatively few ACE-tRNA expression cassettes are required for significant rescue of both CFTR mRNA and channel function from endogenous CFTR genes with CF causing PTCs, which further provides promise for use of ACE-tRNAs as therapeutics for nonsense-associated CF as it reduces the burden of therapeutic delivery efficiency.

### ACE-tRNA expression yields significant rescue of CFTR mRNA expression in G542X-, R1162X- and W1282X-CFTR 16HBEge cells

With the information that as few as 16 copies of ACE-tRNA^Arg^ support significant rescue of CFTR function, we revisited our experiments with transient transfection of plasmids encoding ACE-tRNAs, which do not recover more than 50% of the WT CFTR mRNA. Knowing that the transfection efficiency of 16HBE14o-cells is ∼50%, we wanted to determine if cDNA delivery is impacting our measurements of CFTR mRNA rescue. To account for transfection efficiency, we generated a transfection reporter plasmid containing both an EF1α-driven mNeonGreen (mNG) and 4 copies of either ACE-tRNA^Gly^, ACE-tRNA^Arg^, ACE-tRNA^Leu^, or ACE-tRNA^Trp^ (Fig. 5A). mNG containing constructs permitted us to sort mNG positive (Fig. 5B, green curve) and negative (Fig. 5B, grey curves) cells by fluorescence-activated cell sorting (FACS) 48 hours after transfection and therefore determine ACE-tRNA dependent NMD inhibition in a near-homogeneous population of cells that received ACE-tRNA encoding plasmids. FACS analysis determined the percentage of green cells, those receiving ACE-tRNA plasmid, in fact did not exceed 50% (Fig. 5B, Fig. S4A, Supplemental Table 1). ACE-tRNA mNG positive G542X-, R1162X- and W1282X-CFTR 16HBEge cells exhibited significantly enhanced NMD inhibition over mNG negative cells (Fig. 5C). CFTR mRNA expression for ACE-tRNA^Gly^-G542X was 41%, ACE-tRNA^Arg^-R1162X was 54% and ACE-tRNA^Leu^-W1282X was 87% of WT CFTR. We were excited to see that ACE-tRNA^Leu^ provided strong inhibition of NMD, indicating that when ACE-tRNAs are efficiently delivered to cells, they are exceedingly efficient at inhibiting NMD. The dependence of this result on transfection efficiency also highlights that ACE-tRNA delivery is a significant hurdle to their adoption as a therapeutic (see discussion).

**Figure 5.**
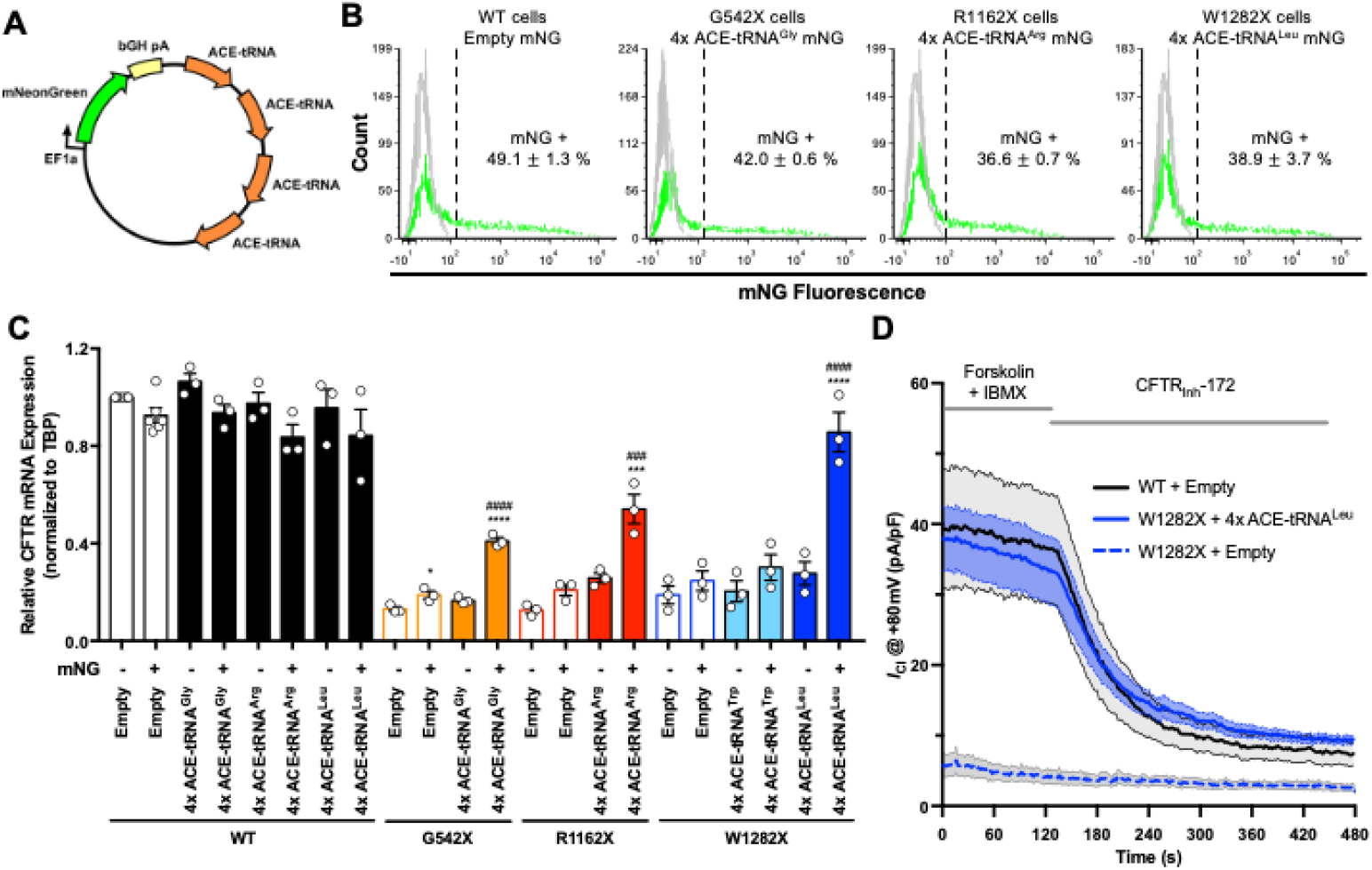
Expression of ACE-tRNAs results in significant and robust increase in CFTR mRNA and channel function. (**A**) Schematic of ACE-tRNA mNG plasmid. (**B**) 48 hrs following transfection of WT, G542X-, R1162X- and W1282X-CFTR cells with plasmids encoding ACE-tRNAs and mNeonGreen (mNG) expression of fluorescent protein was determined by FACS to sort ACE-tRNA transfected cells (mNG positive) from non-transfected cells (mNG negative). Gray curves represent WT cells transfected with empty vectors, which were used to determine the gate (dashed line) for sorting mNG negative (ACE-tRNA not transfected) and positive (ACE-tRNA transfected, green curves) for each cell type and delivered plasmid. The percent mNG positive cell population is indicated in each panel. Representative FACS histograms for WT 16HBE14o-cells (top) and paired empty plasmid transfection of G542X-, W1282X- and R1162X-CFTR cells (bottom) are presented in **Supplemental Fig. 4A**. Representative FACS histograms of cells transfected with control empty plasmids are presented in **Supplemental Fig. 4B**. (**C**) Real-time qRT-PCR quantification of WT, G542X-, R1162X- and W1282X-CFTR mRNA expression in FACS sorted mNG positive and negative cell populations following transfection of mNG plasmids encoding ACE-tRNA^Gly^ (orange bars), ACE-tRNA^Arg^ (red), ACE-tRNA^Trp^ (light blue bars) and ACE-tRNA^Leu^ (dark blue bars). mNG positive and negative WT 16HBE14o-cells transfected with empty (white bars, *n* = 4) and ACE-tRNA^Gly^ (black bars, *n* = 3), ACE-tRNA^Arg^ (black bars, *n* = 3), ACE-tRNA^Trp^ (black bars, *n* = 3), and ACE-tRNA^Leu^ (black bars, *n* = 3), did not exhibit significantly altered CFTR mRNA expression. Transfection of empty mNG plasmids into R1162X- (open red bars, *n* =3) and W1282X-CFTR (open blue bars, *n* =3) cells did not significantly affect CFTR mRNA expression, while mNG positive G542X-CFTR cells (open orange bars, *n* = 3) exhibited a significant but meager increase in CFTR mRNA expression. mNG positive G542X-, R1162X- and W1282X-CFTR cell populations exhibited a significant and robust rescue of CFTR mRNA expression over mNG negative cell populations 48hrs following transfection with 4x ACE-tRNA^Gly^ (solid orange bars, *n* =3), ACE-tRNA^Arg^ (solid red bars, *n* = 3) and ACE-tRNA^Leu^ (solid blue bars, *n* = 3) plasmids, as quantified by real-time qRT-PCR. (**D**) CFTR channel activity was assessed in control empty mNG transfected WT 16HBE14o-cells (black trace, *n* = 7), W1282X-CFTR cells (dashed blue trace, *n* = 5), and 4x ACE-tRNA^Leu^ mNG plasmid transfected W1282X-CFTR cells (blue trace, *n* = 11) using whole-cell voltage clamp. Results in **C** and **D** are presented as average ± SEM, and significance was determined by one-way ANOVA and Tukey’s post-hoc test, where \**P*<0.5 vs. mNG negative, \*\*\*\**P*<0.0001 vs. mNG negative, ^###^*P*<0.001 vs. empty mNG positive and ^####^*P*<0.0001 vs. empty mNG positive.

### Whole-cell patch clamp reveals near-WT rescue of CFTR channel function from endogenous nonsense mutations with ACE-tRNAs

Although we demonstrated significant and robust rescue of CFTR mRNA in G542X-, R1162X- and W1282X-CFTR 16HBEge cells, a rescue of 15-30% of CFTR channel function is ultimately required for a therapeutically relevant outcome (34). To this end, we transfected 4x ACE-tRNA^Leu^ mNG plasmids (Fig. 5A) into W1282X-CFTR 16HBEge cells and utilized the whole-cell patch clamp technique to directly measure CFTR channel function in mNG positive cells 48-72 hours after transfection (Fig. 5D). To fully activate CFTR channels, we pre-incubated cells in forskolin (20μM) and IBMX (100μM) for 5 minutes before starting the recording. We propose that measuring CFTR function under these conditions provides a theoretical upper limit to ACE-tRNA dependent PTC suppression efficiency of endogenous CFTR. Here, ACE-tRNA^Leu^ mNG positive W1282X-CFTR 16HBEge cells exhibited CFTR_Inh_-172 sensitive chloride current magnitudes (Fig. 5D, blue trace, 26.7 ± 4.3 pA/pF at +80mV, n = 11) that were 91% of those measured in WT 16HBE14o-cells (Fig. 5D, black trace, 29.3 ± 6.2 pA/pF at +80mV, n = 7). As expected, W1282X-CFTR 16HBEge cells receiving empty mNG plasmids exhibited no significant CFTR function (Fig. 5D, dashed blue trace, n = 5). These results are consistent with Ussing chamber measurements of R1162X PB 16x ACE-tRNA^Arg^ cells (Fig. 4B), where CFTR channel function appears to closely track CFTR mRNA expression.

## DISCUSSION

Several studies have used nonsense suppressor tRNAs to address PTCs in certain disease-related genes including β-globin in *Xenopus laevis* oocytes (27), xeroderma pigmentosum group A in human cells (28), chloramphenicol acetyl transferase in transgenic mice (40), CDH1 in human cell lines (29), and CFTR in human cells and *Xenopus laevis* oocytes (30). We previously demonstrated using mass spectrometry of the full-length protein product of PTC readthrough that ACE-tRNAs in their current form insert the correct amino acid, providing seamless rescue of the PTC (30). While these studies indicated that ACE-tRNAs are competent to promote suppression of PTCs in mRNA produced from cDNA, they do not fully recapitulate the hurdles seen in the rescue of genuine PTCs in their genomic context. NMD is a eukaryotic translation-dependent mRNA quality control pathway that selectively degrades mRNA transcripts containing PTCs (1, 44). Here we demonstrate CFTR mRNA rescue from 16HBEge cells containing PTCs in the endogenous genomic context. Electrophysiological assays including Ussing chamber short circuit current recording and patch clamp are considered the ‘gold standard’ for evaluating CFTR functional rescue in drug development (32). Using these assays, we demonstrate protein rescue of PTC-containing CFTR, providing proof-of-principle that ACE-tRNAs when delivered as cDNA or RNA can rescue native PTCs. Based on our work here, we note that protein rescue generally tracks with mRNA rescue from NMD. This work establishes a theoretical upper limit to the therapeutic window as >90% of WT function recovered when selecting for efficient delivery of ACE-tRNA as DNA.

While this study represents PTC rescue in one of the most faithful PTC models used to date when assaying for function of ACE-tRNAs, a fundamental limitation is that this is an immortalized cell line and does not fully recapitulate the delivery issues or potential safety issues realized *in vivo*. A notable concern about the safety of ACE-tRNA is that nonsense suppressor tRNAs may cause readthrough of NTCs to toxic effect. In our previous work, we determined via ribosome profiling that readthrough of NTCs is minimal in comparison to PTCs (30). Further, in the context of genetic code expansion, stable expression, and viral delivery of nonsense suppressor tRNAs has been shown to be tolerated in mice (45, 46). Our ability to generate cell lines stably expressing 16 copies of ACE-tRNAs with similar growth rates and morphology to WT parents cells lends further credibility to tolerably low levels of toxicity due to NTC readthrough. It should also be noted that ACE-tRNAs inherently possess a lower level of global NTC readthrough. Unlike the majority of small molecule PTC readthrough agents developed to date, ACE-tRNAs only suppress one of the three stop codons. A further advantage of ACE-tRNAs over small molecule PTC readthrough agents highlighted here is that ACE-tRNAs provide a la carte rescue, that is the amino acid encoded at the PTC can be specified (i.e. Leu at W1282X). While results shown by ourselves and others indicate the safety of ACE-tRNAs, the DNA delivery methods used here require the use of a transfection marker to control for efficient DNA delivery and cannot be directly translated to the clinic. This highlights a remaining roadblock of delivery to the clinical implementation of ACE-tRNAs, however it should be noted that tRNAs as gene therapy cargo possess a number of attractive qualities (31). We have shown here that ACE-tRNAs can be delivered as RNA or DNA, for which there are many therapeutically relevant delivery avenues to pursue.

As demonstrated here, ACE-tRNAs delivered as RNA are competent to rescue the expression of endogenous PTC-containing R1162X-CFTR mRNA. Given that CFTR protein rescue generally tracks with mRNA rescue, it follows that ACE-tRNA delivered as RNA will rescue protein from genes containing endogenous PTCs. However, due to technical limitations of the model cell line used here, we have not demonstrated CFTR protein rescue directly using ACE-tRNA as RNA. While ACE-tRNA delivered as RNA can go through multiple rounds of aminoacylation and translation, it is not a renewable PTC suppression resource like ACE-tRNA delivered as DNA. Therefore, it is likely that repeated delivery of ACE-tRNA would be needed to sustain a therapeutic level of target protein expression and that modifications to the RNA may be necessary to extend the half-life of the ACE-tRNA therapeutic agent. However, because ACE-tRNAs are only ∼76 nucleotides in length, many of the existing mRNA and short RNA delivery technologies can likely be adapted for delivery of ACE-tRNAs. These delivery methods include but are not limited to cationic or ionizable lipids and lipid-like materials and polymers/nanoparticles (47). In contrast to ACE-tRNA delivered as RNA, delivery as DNA likely will result in augmented persistence of activity.

When ACE-tRNAs are efficiently delivered as DNA, we demonstrate here rescue of both PTC-containing CFTR mRNA and near 100% rescue of CFTR protein, far exceeding the therapeutic threshold for functional CFTR rescue of 15-30% (34). Due to the small size of ACE-tRNA expression cassette (∼250 nucleotides), there are a number of avenues available to deliver ACE-tRNA DNA expression cassettes. The small size of the ACE-tRNA expression cassette allows for optimization of cassette copy number with a minimal vector size which in turn has been shown to increase intracellular mobility, nuclear entry, and gene therapy agent expression (48-50). Naked DNA delivery is an attractive gene therapy approach as it has inherently low immunogenicity, can be designed to remain episomal or integrate, and can be paired with several technologies including nanoparticle formulations, liposomes, pressure injection, and electroporation (51-53). When plasmid DNA is made resistant to silencing, several studies have shown extended expression (months) in murine lung following plasmid DNA delivery by polyethyleneimine/DNA complexes (54), cationic liposome/DNA complexes (55), and DNA electroporation (56).

The small expression cassette size of ACE-tRNAs also make them amenable to delivery by a variety of viral vectors. Viral delivery is an attractive gene therapy approach as viral cargo delivery is usually much more efficient than non-viral delivery and has the advantage of cell-type-specific transduction. There are a number of candidate viral vectors, some of which include adenovirus, adeno-associated virus (AAV), and lentivirus (57). Recently the FDA approved the use of the first AAV therapeutic for total gene replacement therapy of *RPE65* (retinal pigment epithelial-specific 65 kDa protein) for treatment of the inherited retinal disorder Leber congenital amaurosis type-2 (Luxturna®). This demonstrates the possibility of therapeutic a possible route to viral delivery of ACE-tRNAs. Of note, ACE-tRNAs are particularly amenable to AAVs as their small size allows for multiple expression cassettes despite the relatively small cargo capacity of AAVs (<4.7 kb). Indeed, for many diseases associated with nonsense mutations, AAV total gene replacement is not viable due to the large size of the gene in question. In contrast, ACE-tRNA-mediated PTC suppression is independent of target gene size, meaning ACE-tRNAs delivered as an AAV can rescue gene targets of any size. As we demonstrated here that integrating up to 16 copies of ACE-tRNA^Arg^ was well tolerated in 16HBEge cells, the use of integrating lentiviral vectors for one-time delivery of ACE-tRNAs is another attractive approach.

In summary, nonsense mutations cause ∼11% of all serious genetic diseases including CF, for which there are currently few therapeutic options even given their unitary mechanism. Our results suggest that nonsense suppressor ACE-tRNAs are capable of rescuing both PTC-containing mRNA from NMD and functional protein expression. Coupled with an efficient RNA or DNA delivery technology, ACE-tRNAs may provide relief for a variety of genetic diseases in the future.

## MATERIALS AND METHODS

### Materials for Molecular Biology

Unless otherwise specified, all molecular biology reagents were obtained from New England Biolabs. For all PCR reactions Q5 High-Fidelity 2X Master Mix (NEB M0492L) was used, Gibson assembly was performed using NEBuilder HiFi DNA Assembly Master Mix (NEB E2621L), Golden Gate reactions used T4 DNA ligase (NEB M0202L) and restriction enzymes from NEB, gel purifications were performed using the Monarch DNA Gel Extraction Kit (NEB T1020L), and minipreps were performed using the Monarch Plasmid Miniprep Kit (NEB T1010L). NEB 5-alpha Competent *E. coli* (NEB C2987H) were used for all bacterial transformations unless otherwise noted. All oligonucleotides were synthesized by Integrated DNA Technologies. All cDNA preparations were performed using the NucleoBond Xtra Midi EF kit (Macherey Nagel Ref 740420.50).

### DNA constructs/Plasmid generation

The *piggyBac* transposon plasmid (PB plasmid) with *piggyBac* inverted terminal repeats (ITRs) flanking an ampicillin resistance marker, pBR322 origin, short ubiquitin C promoter (UbC)-puromycin *N*-acetyltransferase-SV40 pA, and EcoRV, PmlI, PmeI, SmaI multiple cloning site in pUC57 was synthesized (Genscript). The hyperPB transposase sequence (58) was synthesized as gBlocks (IDT, Coralville, IA, USA) and Gibson Assembled into pcDNA3.1(+). For further specifics of plasmid cloning methods see supplemental materials and methods.

### Cell culture

Parental 16HBE14o-human bronchial epithelial cells, which were generated by Dieter Gruenert (University of California, San Francisco) (59), and 16HBEge cells with engineered CFTR variants, CFF-16HBEge CFTR G542X, CFF-16HBEge CFTR R1162X and CFF-16HBEge CFTR W1282X cells, were obtained from the Cystic Fibrosis Foundation Therapeutics Lab (Lexington, MA, USA) (32). HBE cells were cultured in Minimum Essential Medium (MEM; Gibco, #11095-080) supplemented with 10% Fetal Bovine Serum (Gibco, #26140-079) and 1% Penicillin/Streptomycin/Glutamine (Gibco, #10378-016) in a 37 °C/5% CO_2_ humidified incubator. Plates were coated with coating solution [LHC-8 basal medium (Gibco, #12677-027), 1.34 µl/ml Bovine serum albumin 7.5% (Gibco, #15260-037), 10 µl/ml Bovine collagen solution, Type 1 (Advanced BioMatrix, #5005-100ML) and 10 µl/ml Fibronectin from human plasma (Thermo Fisher Scientific, #33016-015)] for 2-3 h at 37 °C/5% CO_2_ and stored at 4 °C after thorough removal of coating solution.

### Transfection and Nucleofection of HBE cells

HBE cells were transfected with Lipofectamine LTX and Plus reagent (Invitrogen by Thermo Fisher Scientific, #15338-100) according to manufacturer’s guidelines with slight modifications. Briefly, on the day of transfection, 500,000 cells per well were plated in a 6-well plate and incubated in a cell culture incubator for 20-30 minutes. 2.5 µg of DNA and 2.5 µl of PLUS reagent were diluted in 125 µl of Opti-MEM (Gibco, #51985-034), and 5 µl of Lipofectamine LTX reagent was diluted in Opti-MEM. Diluted cDNA and Lipofectamine LTX reagent were combined, mixed thoroughly and incubated for 5 minutes at room temperature. The transfection mixture was then added dropwise to pre-plated cells. The media containing transfection mix was replaced with normal cell culture media 16-24 hours post transfection. HBE cells were nucleofected with the Amaxa 4D-Nucleofector system (Lonza, Basel, Switzerland) using the Amaxa SG cell line 4D-Nucleofector X Kit S (Lonza, #V4XC-3032). Each well of a black 96-well plate was filled with 95 µl of cell culture media and pre-incubated in a humidified 37 °C/5% CO_2_ incubator, and an aliquot of cell culture media was pre-warmed in a 37 °C water bath. For each reaction, 200,000 cells were centrifuged at 90 x g for 10 minutes, resuspended in 20 µl of supplemented SG 4D-Nucleofector X solution and mixed with 1 µg of RNA (maximum 10% of final sample volume). A total volume of 20 µl was transferred to a well of Nucleocuvette Strip. The nucleofection process was started by placing the strip into the 4D-Nucleofector X Unit and running the CM-137 Nucleofector program from the 4D-Nucleofector Core Unit. Cells were incubated at room temperature for 10 minutes and resuspended with 80 µl of pre-warmed cell culture media. 15 µl of nucleofected cells were plated in pre-incubated cell culture plate and maintained in a humidified 37 °C/5% CO_2_ incubator until analysis.

### Stable cell line generation

To generate stable HBE cell lines, the *piggyBac* (PB) transposon vector expressing gene of interest (NLuc PTC reporter or ACE-tRNA) was co-transfected with the PB transposase vector. For each well of a 12-well plate, 250,000 cells were seeded on the day of transfection and transfected with 0.5 µg of PB transposon vector and 0.5 µg of empty vector as a negative control or 0.5 µg of PB transposon vector and 0.5 µg of PB transposase vector using Lipofectamine LTX and Plus reagent as described above. Stable cells were selected with 0.5 µg/ml of puromycin (InvivoGen, #ant-pr-1) 1-2 days post transfection until no viable cells were detected in the negative control condition.

### Fluorescence-activated cell sorting (FACS)

For fluorescence-activated cell sorting (FACS) experiment, cells were washed with Dulbecco’s phosphate-buffered saline (DPBS; Gibco, #14190-144) twice, dissociated with TrypLE Express (Gibco, #12604-013), pelleted at 1000 rpm (∼500 x g) for 5 minutes, resuspended with DPBS with calcium and magnesium (Gibco, #14040-133) and strained with 40 µm Cell Strainer. Cells were then sorted using the BD FACSAria II cell sorter (BD Biosciences), and data were analyzed by using FCS Express 7 flow cytometry software (De Novo Software, Pasadena, CA, USA).

In order to sort ACE-tRNA expressing cells from ACE-tRNA not expressing cells, HBE cells were transfected with the plasmid containing ACE-tRNA and mNeonGreen (mNG) fluorescent protein in one plasmid. Approximately 36-48 hours after transfection, cells were prepared as described above for FACS experiment. Empty vector expressing cells were used to determine the gate for mNG negative and positive cells. Each population of mNG negative and positive cells was then sorted into a 15 ml conical tube using the BD FACSAria II cell sorter. Total RNA was isolated from sorted cells and used for RT-qPCR as described below.

### Luciferase reporter assay

PTC readthrough by ACE-tRNA was assessed by measuring NanoLuc (NLuc) luciferase activity using the Nano-Glo Luciferase Assay System (Promega, #N1130) on a Synergy2 multi-mode microplate reader (BioTek Instruments, Winooski, VT, USA). Using Lipofectamine LTX and Plus reagent as described above, HBE cells were co-transfected with NLuc, NLuc-PEST or Sec-NLuc PTC reporters and ACE-tRNA, or HBE cells stably expressing PTC reporters were transfected with ACE-tRNA. In each well of a black 96-well plate, 15 µl of DPBS was added for cells expressing NLuc and NLuc-PEST PTC reporters, whereas 15 µl of media from transfected cells were transferred for cells expressing Sec-NLuc PTC reporter. Then, 15 µl of reconstituted Nano-Glo Luciferase Assay Reagent (1:50 of Nano-Glo Luciferase Assay Substrate to Nano-Glo Luciferase Assay Buffer) was added to each well for luminescence recording using a Synergy2 multi-mode microplate reader. Each condition was assayed in triplicate, and cells expressing empty vector instead of ACE-tRNA were included as a negative control. Luciferase activity was displayed as raw NLuc luminescence or normalized NLuc luminescence to background luminescence.

### Quantitative reverse transcription polymerase chain reaction (RT-qPCR)

Total RNA was isolated from cells with the Monarch Total RNA Miniprep Kit (New England BioLabs, #T2010S) according to manufacturer’s recommendations. RNA quantity and quality were measured with a NanoDrop One^c^ Spectrophotometer (Thermo Fisher Scientific, Waltham, MA, USA). Subsequently, one-step reverse transcriptase and quantitative PCR (RT-qPCR) was performed on the QuantStudio 3 Real-Time PCR System (Applied Biosystems, Waltham, MA, USA) using the Luna Universal One-Step RT-qPCR Kit (New England BioLabs, #E3005L or #E3006E) according to manufacturer’s instructions and then analyzed with QuantStudio Design & Analysis Software v1.5.1 (Applied Biosystems, Waltham, MA, USA). Briefly, the reaction mixture contained 10 µl 2x Luna Universal Probe One-Step Reaction Mix, 1 µl 20x Luna WarmStart RT Enzyme Mix, 1 µl 20x CFTR-specific primers and probe, 1 µl TBP-specific primers and probe, 100-500 ng of template RNA, and nuclease-free water in a final volume of 20 µl. The amplification conditions were as follows: one cycle of 10 min at 55 °C and one cycle of 3 min at 95 °C followed by 40 cycles of 15 s at 95 °C and 30 s at 60 °C. Fluorescence was detected at the end of the 60 °C extension step. Each sample was quantified in triplicate, and a no-template control (NTC) reaction, with just nuclease-free water, was included as a negative control. The fold difference in CFTR gene expression normalized to TBP was calculated using the comparative Ct method, 2^-ΔΔCt^. The following target-specific primers and probe used in RT-qPCR experiments were purchased from Integrated DNA technologies (Coralville, IA, USA): CFTR probe, CFTR primer 1, and CFTR primer 2 and endogenous control TBP-specific TBP probe, TBP primer 1, and TBP primer 2 (sequences in Supplementary Table 2). The final concentration of primers was 500 nM each and probe was 250 nM.

### Ussing chamber studies of HBE cells

HBE cells were seeded onto 6.5mm Transwell Permeable Support with 0.4 µm Polycarbonate Membrane Inserts (Corning, #3413) pre-coated with Collagen from rat tail (Sigma-Aldrich, #C7661-25MG) 5-6 days before assay and maintained in normal cell culture media at 37 °C and 5% CO_2_. Cells were grown until the formation of electrically tight epithelial monolayer was confirmed by measuring transepithelial resistance with EVOM^2^ Epithelial Voltohmmeter (World Precision Instruments, Sarasota, FL, USA). Short circuit current (*I*_sc_) was measured under voltage clamp condition to determine CFTR channel function. Cell monolayers were mounted in Ussing chambers (Physiologic Instruments, San Diego, CA, USA) filled with Ringer’s solution (in mM: 135 NaCl, 5 HEPES, 0.6 KH_2_PO_4_, 2.4 K_2_HPO_4_, 1.2 MgCl_2_, 1.2 CaCl_2_, 5 D-Glucose adjusted to pH 7.4) at 37 °C and gassed with compressed air. After baseline recordings, cells were treated from the apical side with 100 µM amiloride to block epithelial sodium channels and then with 100 µM 4,4’-diisothiocyanostilbene-2,2’-disulfonic acid (DIDS) to block Ca^2+^-activated chloride channels. Then, the solution in the apical side was replaced with a low Cl^-^ solution (in mM: 135 Na-Gluconate, 5 HEPES, 0.6 KH_2_PO_4_, 2.4 K_2_HPO_4_, 1.2 MgCl_2_, 1.2 CaCl_2_, 5 D-Glucose adjusted to pH 7.4). CFTR-mediated Cl^-^ current was measured by adding CFTR activators, 10 µM forskolin and 100 µM 3-isobutyl-1-methlyxanthine (IBMX), and CFTR inhibitor, 20 µM CFTR inhibitor-172 (CFTR_Inh_-172), to the apical side. Data were acquired with Acquire & Analyze 2.3 software (Physiologic Instruments, San Diego, CA, USA). The maximum change in I_sc_ in response to additions of CFTR activators and CFTR inhibitor were quantified as measures of functional CFTR Cl^-^ transport activity.

### Macroscopic Recordings of CFTR Currents

Cells were transfected using Lipofectamine™ LTX Reagent with PLUS™ Reagent (Invitrogen) with a modified reverse transfection as described above. W1282X-CFTR 16HBEge cells were transfected with either a plasmid containing 4 copies of ACE-tRNA^Leu^ with mNG or empty plasmid encoding only mNG. Results were compared to WT 16HBE14o-cells transfected with empty mNG plasmid. After transfection, cells were lifted using Versene (Gibco, #15040-066), plated on 35mm dishes coated with human fibronectin (as previously described) at low density, and cells in isolation were patched 2-3 days post transfection. Cells were continuously perfused with external solution (in mM: 145 TEA-Cl, 10 CaCl_2_, 10 HEPES adjusted to pH 7.4 with TEA-OH) supplemented with 10 uM Forskolin (Sigma) and 100 uM IBMX (Sigma) for maximal activation of CFTRs; dishes of cells were incubated in this solution for at least 5 minutes prior to recording. Green cells were patched using thick-walled borosilicate glass pipettes (Sutter, Novato, CA, USA) pulled to approximately 7 ± 3 MOhm using a vertical puller (PC-100, Narishige, Amityville, NY, USA). Pipettes were filled with internal solution (in mM: 30 CsCl, 110 Cs-aspartate, 2.5 CaCl_2_, 2.5 MgCl_2_, 10 HEPES, 5 CsEGTA adjusted to pH 7.4 with CsOH). Internal and external solutions were formulated to eliminate non-chloride contaminating currents (i.e., TEA and cesium to block potassium currents and no sodium present to eliminate sodium currents) and feature a reversal potential for chloride of approximately -36 mV. Currents were amplified using an Axon 200B amplifier (Axon Instruments, Hamden, CT, USA), sampled at 10 kHz and filtered at 2 kHz using a Digidata 1440A, and recorded using pClamp 10 (Molecular Devices, San Jose, CA, USA). Whole-cell capacitance and series resistance were compensated (approximately 70%) and a linear ramp voltage protocol from -80 mV to +80mV (200ms) was then applied from the holding potential (™40mV) to collect current traces at 0.5 Hz for 60 sweeps. Perfused solutions were then rapidly exchanged using a fast-step perfusion barrel (VC-8, Warner Instruments) to an external solution containing 20 µM CFTR_Inh_-172 to block CFTR currents and the ramp voltage protocol was continued at 0.5 Hz for 180 more sweeps. Recordings were analyzed using Clampfit 10, whereby peak outward current near +80 mV was divided by cell capacitance, as determined by the membrane test function in pClamp, to yield current densities (pA/pF). Peak outward current density at +80 mV was then plotted versus time using Prism 9 (Graphpad) for comparison of treatment groups and controls. Only experiments where a gigaohm seal was achieved before break-in and had a measured reversal potential that fell within 7 mV of -36 mV were used in analysis.

### Quantification of stably integrated copy number using quantitative real-time PCR

Genomic ACE-tRNA integration of the cell population was quantified by performing quantitative real-time PCR (qPCR). Genomic DNA (gDNA) was extracted using the Monarch Genomic DNA Purification Kit (New England BioLabs, #T3010S) according to manufacturer’s instructions. Genomic integrants of the PB transposon expressing 16 copies of ACE-tRNA^Arg^ cassette (PB 16x ACE-tRNA^Arg^) were detected by targeting either the puromycin cassette *(puro)* or PB backbone (PB backbone) from the PB transposon vector. Puro-specific *puro* probe, *puro* primer 1, and *puro* primer 2 and PB backbone specific PB backbone probe, PB backbone primer 1, and PB backbone primer 2 were purchased from Integrated DNA technologies (Coralville, IA, USA). The gDNA content was normalized based on the amplification of the *Ribonuclease P* (*RNase P*) gene using *RNase P* probe purchased from Applied Biosystems and *RNase P* primer 1 and *RNase P* primer 2 purchased from Invitrogen. The final concentration of primers was 500 nM each and probe was 250 nM. Real-time PCR was performed on the QuantStudio 3 Real-Time PCR System using the PrimeTime Gene Expression Master Mix (Integrated DNA Technologies, #1055772) according to manufacturer’s guidelines and then analyzed with QuantStudio Design & Analysis Software v1.5.1. Briefly, the reaction mixture contained 10 µl 2x PrimeTime Gene Expression Master Mix, 1 µl 20x *puro*- or PB backbone-specific primers and probe (target gene), 1 µl *RNase P*-specific primers and probe (reference gene), DNA template and nuclease-free water in a final volume of 20 µl. PCR cycling conditions were one cycle of 3 min at 95°C followed by 40 cycles of 15 s denaturation at 95°C and 1 min annealing and extension at 60°C. A standard curve was constructed by serial dilutions of a PB 16x ACE-tRNA^Arg^ plasmid with *puro*- or PB backbone-specific primers and probe and Hs_RPP30 Positive Control (Integrated DNA technologies, #10006622) with *RNase P*-specific primers and probe. The number of ACE-tRNA^Arg^ copies in 10 ng or 1 ng DNA from cells stably integrated with ACE-tRNA^Arg^ was calculated by plotting cycle threshold (C_t_) to the linear equation from the standard curve. All qPCR reactions were run in triplicate. A NTC reaction with just nuclease-free water was included as a negative control, and 10 ng or 1ng DNA from cells not integrated with ACE-tRNA^Arg^ was assayed to confirm specificity of *puro*- and PB backbone-specific primers and probe.

### Statistical analysis

Data are presented as mean ± standard error of the mean (SEM). GraphPad Prism 7 software (GraphPad Software, San Diego, CA, USA) was used for all statistical analysis. Comparisons between two groups were performed using an unpaired Student’s t-test. Multiple comparisons were analyzed using a one-way analysis of variance (ANOVA) followed by Tukey’s multiple comparisons test. *p* values < 0.05 were considered statistically significant.

## Acknowledgments

We thank members of the Lueck Laboratory and Dr. Amy M. Martin for reading and editing the manuscript and constructive discussion throughout the study. We thank T.N. Lueck with help generating figures. We would like to thank the Cystic Fibrosis Foundation Therapeutics Lab and Dr. Hillary Valley for providing 16HBE14o- and 16HBE14ge cell lines used in this study and Dr. David MacLean for extended use of his electrophysiology hardware.

## Author Contributions

J.J.P., W.K., and J.D.L designed the study. J.J.P., W.K., M.T.S, K.M.E and J.D.L. performed experiments. J.J.P., W.K., M.T.S, and J.D.L. analyzed the data, and constructed the figures. J.J.P, W.K., and J.D.L. wrote the manuscript. All authors read and revised the manuscript.

## Data and Material Availability

All data are available in the main text or the supplementary materials.

