## Supplemental Information for "Efficient Suppression of Endogenous CFTR Nonsense Mutations Using Anticodon Engineered Transfer RNAs"

**Supplementary Materials**

**Materials and Methods**

**Plasmid DNA cloning methods**

The human EF1a promoter including intron A was PCR amplified from pEF-GFP (a gift from Connie Cepko, Addgene plasmid #11154; <http://n2t.net/addgene:11154> ; RRID:Addgene 11154) (*60*) using primers EF1a pcDNA gibF and EF1a pcDNA gibF (Supplementary Table 2). A pcDNA3.1(+) plasmid containing EGFP was PCR amplified using primers pcDNA prom gibF and pcDNA prom gibR. The resulting DNA fragments were resolved on a 0.6% TAE agarose (Fisher Scientific) gel containing ethidium bromide and gel purified from excised bands as per the Monarch DNA gel Extraction Kit. The DNA fragments were assembled using the NEBuilder HiFi mix with reactions containing 30 fmol of each insert and 10 fmol of plasmid backbone DNA. The resulting pcEF1a EGFP plasmid was transformed into NEB 5-alpha competent cells, miniprepped using the Monarch Plasmid miniprep kit, and sequence verified by Sanger sequencing (Eurofins Genomics). The EF1a-EGFP-bGH cassette was then transferred to the PmeI site of the PB plasmid for stable genomic integration via PiggyBac transposase. The EF1a-EGFP-HA-bGH cassette was amplified with PROMgaF -> PmeI PB and bGHgaR -> PmeI PB2, the PB plasmid was restriction enzyme digested with PmeI, both DNA fragments were gel purified, assembled via Gibson assembly, transformed, miniprepped, and sequence verified as outlined above.

To allow for stable integration and expression of the NanoLuc PTC reporters they were cloned into the PB plasmid EF1a plasmid. EF1a-EGFP-bGH PB was restriction enzyme digested with EcoRI-HF and XbaI and gel purified as outlined above. NanoLuc TAA, NanoLuc TAG, and NanoLuc TGA were PCR amplified from the respective pcDNA3.1(+) plasmids using primers NLuc Bare/PEST -> EF1a gibF and NLuc Bare/Sec -> EF1a gibR. NanoLuc PEST TAA, NanoLuc PEST TAG, and NanoLuc PEST TGA were PCR amplified from the respective pcDNA3.1(+) plasmids using primers NLuc Bare/PEST -> EF1a gibF and NLuc PEST -> EF1a gibR. NanoLuc Sec TAA, NanoLuc Sec TAG, and NanoLuc Sec TGA were PCR amplified from the respective pcDNA3.1(+) plasmids using primers NLuc Sec -> EF1a gibF and NLuc Bare/Sec -> EF1a gibR. All PCR products were gel purified, assembled into the EF1a PB plasmid by Gibson assembly, transformed, miniprepped, and sequence verified as outlined above.

For efficient expression of multiple ACE-tRNA copies from a single plasmid we reasoned that optimal transcription requires adequate spacing between ACE-tRNA cassettes. To that end we had a plasmid synthesized containing 4 ACE-tRNA cloning sites containing a pair of BbsI, BsaI, BsmBI, or BtgZI restriction enzyme sites flanked by a tRNA^Tyr^ 55 bp leader sequence (*30*) and poly-T RNA polymerase III terminator in pUC57 mini Amp (with the BsaI restriction enzyme site removed from the backbone). Each ACE-tRNA was ordered as a pair of single stranded DNA oligos (Supplementary Table 2), which when annealed leave overhangs corresponding to the overhangs for Golden Gate assembly into the 4x ACE-tRNA plasmids. Each forward and reverse oligo pair was annealed at 100 ng/μL in IDT annealing buffer (100 mM potassium acetate, 30 mM HEPES, pH 7.5) by heating to 95 °C for 10 minutes followed by slow cooling to 4 °C over 45 minutes. The annealed oligos were diluted to 10 ng/μL in molecular biology grade H_2_O (GE Lifesciences SH30538.03) before being used for Golden Gate assembly. Each annealed ACE-tRNA oligo pair was cloned sequentially into the 4x tRNA plasmid by Golden Gate assembly using each of the restriction enzymes in turn (BbsI-HF (NEB R3539L), BsaI-HFv2 (NEB R3733L), BsmBI (NEB R0580), BtgZI (NEB R0703)). A typical Golden Gate reaction mix contained 4.5 μL molecular biology grade H_2_O, 1 μL 10 CutSmart Buffer (NEB B7204S), 1 μL 10 mM ATP (NEB P0756L), 1 μL (400 units) T4 DNA ligase (NEB M0202L), 0.5 μL restriction enzyme, 1 μL (10 ng) annealed tRNA oligos, and 1 μL (100 ng) 4x tRNA GG destination plasmid. For reactions with BsmBI, 10x NEBuffer 3.1 (NEB B7203S) was substituted for 10x Cutsmart buffer. Following assembly of the above Golden Gate reaction components on ice the sample were transferred to a thermocycler and subjected to the following cycle, 37 °C (5 minutes), 20 °C (5 minutes), for 30 cycles each, followed by 37 °C (10 minutes), 80 °C (10 minutes), for 1 cycle each. Following the Golden Gate assembly, 1.5 μL of the reaction was transformed, miniprepped, and sequence verified as outlined above.

Due to the difficulties inherent in assembling plasmids with many repeats we employed a modular system for assembly of the 16x ACE-tRNA cassettes. As this system is a derivative of the pMuLE (*61*) based modular lentiviral expression system we termed it pDonkey. The pDonkey destination cloning site consists of a polycistronic lacUV5 driven chloramphenicol acetyl transferase/*CcdB* (*E. coli* positive/negative selection markers respectively) cassette flanked by Gateway attR1 and attR2 sites. The pDonkey cloning site was cloned into the PB plasmid by amplifying pMuLE Dest (pMuLE Lenti Dest eGFP was a gift from Ian Frew (Addgene plasmid #62175 ; <http://n2t.net/addgene:62175> ; RRID:Addgene_62175)) (*61*) with primers EcoRVattR1/R2GAF and EcoRVattR1/R2GAR. The PB plasmid was restriction enzyme digested with EcoRV and gel purified as outlined above. The pDonkey cassette was cloned into the PB plasmid by Gibson assembly, transformed into One Shot ccdB Survival 2 T1^R^ Competent Cells (Thermo Scientific Invitrogen A10460), miniprepped, and sequence verified as above. Four different entry clones were used for each 16x ACE-tRNA clone assembled which we termed Piece 1, Piece 2, Piece 3, and Piece 4. The cloning cassettes for each Piece are flanked by lambda phage att sites for Gateway assembly. The Piece 1 tRNA expression cassette is flanked by attL1 and attR5, Piece 2 by attL5 and attL4, Piece 3 by attR4 and attR3, and Piece 4 by attL3 and attL2. The tRNAs for each 4x ACE-tRNA plasmid were cloned as outlined above into Pieces 1-4. After each of the 4x ACE-tRNA Piece 1-4 and the PB Donkey were constructed and sequence verified, the final 16x ACE-tRNA clones were assembled as per the LR Clonase II Plus enzyme mix instructions (Thermo Scientific Invitrogen 12538120). Briefly, each of the entry vectors (Pieces 1-4) were diluted to 10 fmol/μL (for Pieces 1-4 this is 22 ng/μL) and the PB Donkey destination vector was diluted to 20 fmol/μL (for the PB Donkey this is 80 ng/μL). A typical reaction was 1 μL of each entry clone at the above concentration, 1 μL of destination clone at the above concentration, 3 μL TE (1X solution, pH 7.5, molecular biology grade, ultrapure, Thermo Scientific), and 2 μL LR Clonase II Plus enzyme mix (Thermo Scientific Invitrogen 12538120). The reaction was held at 25 °C in a thermocycler for 24 hours and 5 μL of the reaction was transformed into 50 μL of NEB5a chemically competent *E. coli* (NEB C2987H). The cells were plated on Miller LB Agar (Fisher Bioreagents) containing 50 μg/mL kanamycin (Fisher Scientific) and grown overnight at 37 °C. Colonies were inoculated into ZYM-505 media containing 50 μg/mL kanamycin and grown overnight at 37 °C shaking at 250 rpm. Following minipreps the clones were screened for successful assembly by restriction enzyme digest and the 16x ACE-tRNA cassette was sequenced as above.

To select for cell transfected with ACE-tRNAs, we cloned an EF1a-mNeonGreen-bGH cassette into the vector backbone of Piece 3 4xACE-tRNA. Each Piece 3 4xACE-tRNA was restriction enzyme digested with ZraI and gel purified as outlined above. The EF1a-mNeonGreen-bGH cassette was amplified with primers EF1a P3 ZraI gibF and EF1a P3 ZraI gibR and gel purified as outlined above. The Piece 3 4xACE-tRNA EF1a-mNeonGreen-bGH was Gibson assembled, transformed, miniprepped, and sequence verified as outlined above.

**Supplemental Figures**

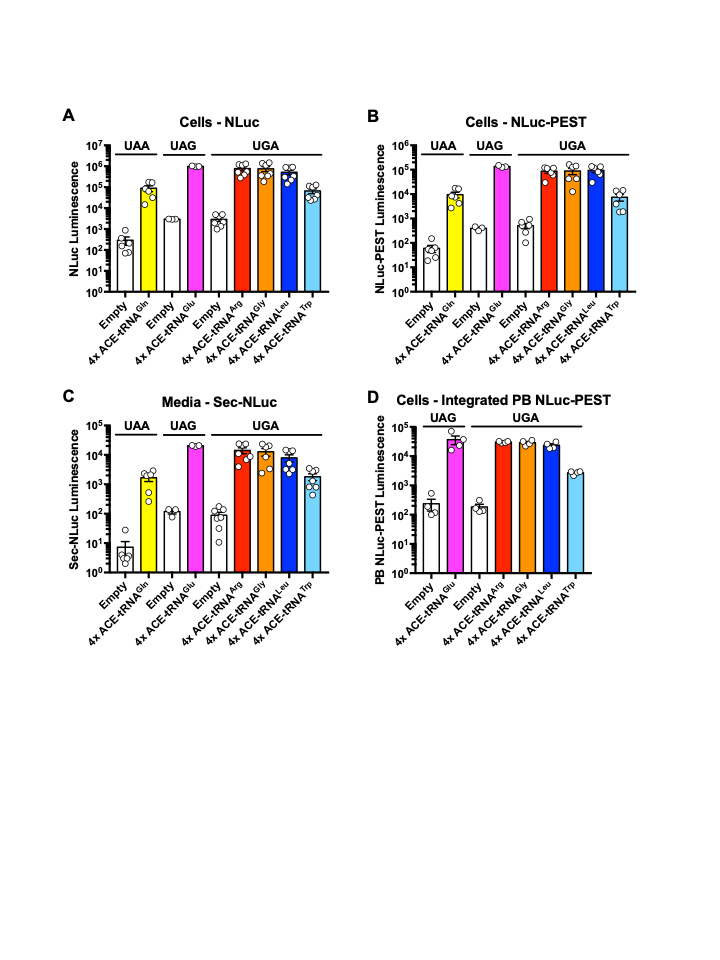

**Figure S1. (Raw luminescent values of Fig. 1) ACE-tRNAs are efficient at suppressing PTCs encoded in NLuc reporters in 16HBE14o- cells.** (**A-C**) Raw NLuc luminescence following co-transfection of ACE-tRNA plasmids and (**A**) NLuc PTC, (**B**) NLuc-PEST PTC, or (**C**) Sec-NLuc PTC reporter plasmids in WT 16HBE14o- cells. ACE-tRNAs efficiently rescue luminescence of three NLuc PTC reporters with varying degrees compared to empty vector control. (**D**) Stably integrated NLuc-PEST luminescence PTC construct reports on ACE-tRNA PTC suppression activity. Data are presented in Log 10 scale of raw luminescence ± SEM. *n* = 3-7, with each replicate is represented as a circle.

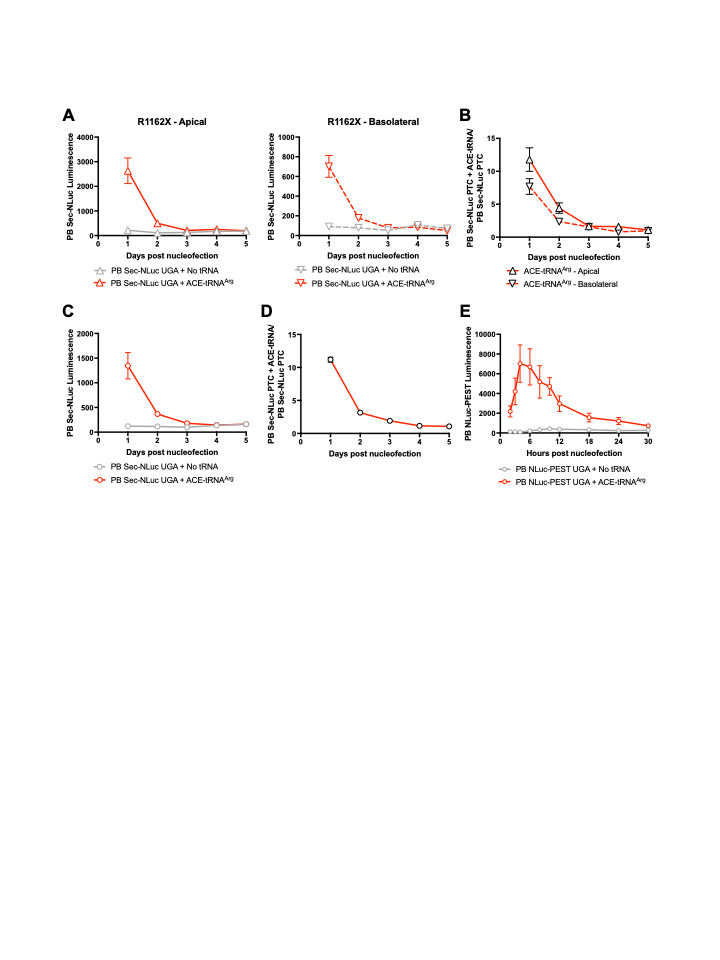

**Figure S2. Comparison of Sec-NLuc UGA and PEST-NLuc UGA constructs to report time-course of ACE-tRNA^Arg^ RNA suppression activity.** (**A-D**) R1162X PB Sec-NLuc UGA cells were nucleofected with and without ACE-tRNA^Arg^ delivered as RNA and seeded onto (**A-B**) 6.5 mm Transwell or (**C-D**) 96-well plate. Nonsense suppression mediated by ACE-tRNA^Arg^ RNA was detected by luminescence measurement of secreted NLuc containing media for 5 consecutive days post nucleofection from apical (solid line) and basolateral (dashed) sides of Transwell inserts. Results are presented as (**A, C**) raw luminescence and (**B, D**) normalized luminescence (ACE-tRNA^Arg^ RNA/ No tRNA), the PTC suppression activity of ACE-tRNA^Arg^ as RNA significantly decreases in 48hrs post nucleofection. (**E**) Raw luminescence measurements reported in **Fig 3A** for R1162X PB PEST-NLuc UGA cells nucleofected with (red) and without (gray) ACE-tRNA^Arg^ in RNA. Data are presented as average ± SEM. Each symbol represents *n* = 3.

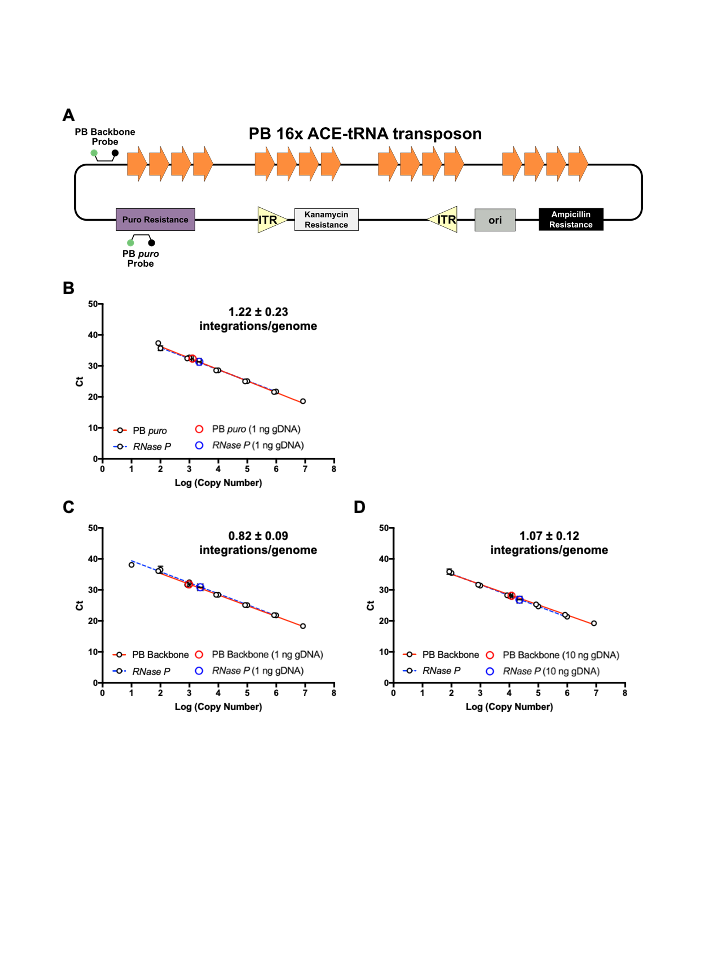

**Figure S3. ACE-tRNA^Arg^ expression cassettes are stably integrated in R1162X-CFTR cells using PB 16x ACE-tRNA transposon.** (**A**) Schematic of PB 16x ACE-tRNA transposon cDNA with location of qPCR probes for puro transgene and backbone (**B-D**) qPCR standard curves generated from 10-fold dilutions of 16x ACE-tRNA^Arg^ (red lines) and RNase P (blue dashed lines) cDNA. qPCR quantification of *puro* transgene (red circle) and endogenous *RNase P* gene (blue circle) from gDNA isolated from R1162X PB 16x ACE-tRNA^Arg^ cells. (**B**) 1 ng of input gDNA extracted from R1162X PB 16x ACE-tRNA^Arg^ with qPCR measurement of PB *puro* specific primers and probe (red circle) and *RNase P*-specific primers and probe (blue circle). (**C**) 1 ng of input gDNA extracted from R1162X PB 16x ACE-tRNA^Arg^ with qPCR quantification of PB backbone specific primers and probe (red circle) and *RNase P*-specific primers and probe (blue circle). (**D**) 10 ng of input gDNA extracted from R1162X PB 16x ACE-tRNA^Arg^ with qPCR quantification of PB backbone specific primers and probe (red circle) and *RNase P*-specific primers and probe (blue circle). Number of genomic PB integration events were calculated as described in **Fig. 3G** and added as inserts to each figure panel. Data are presented as average ± SEM. Each symbol represents *n* = 3.

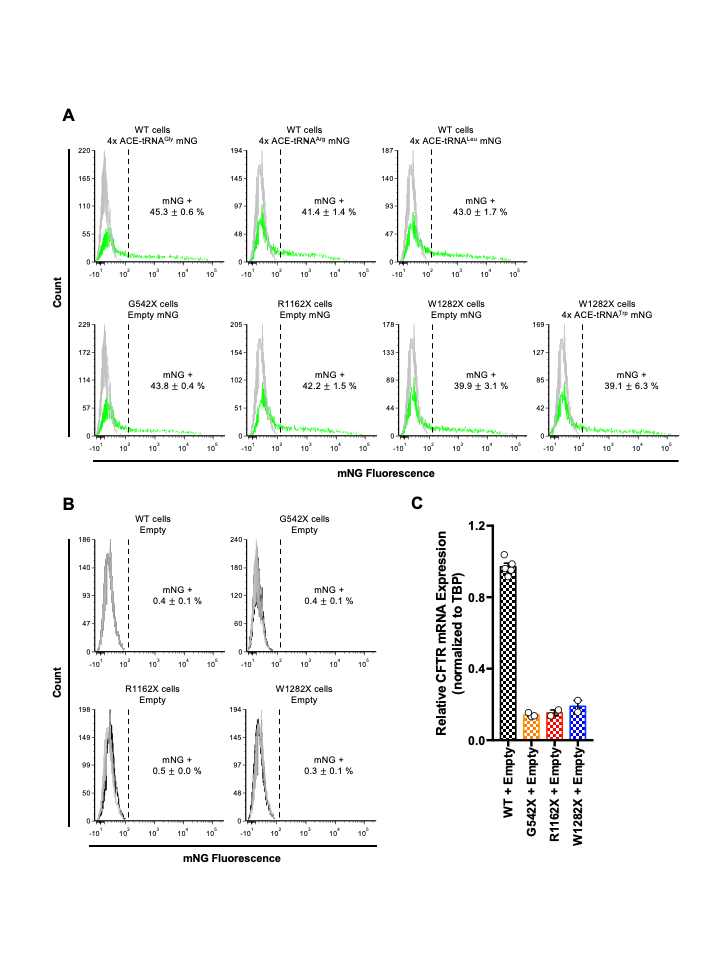

**Figure S4. FACS analysis of cells transfected with control plasmids.** (**A**) Representative FACS analysis histograms of WT 16HBE14o- cells transfected with 4x ACE-tRNA^Gly^, 4x ACE-tRNA^Arg^ and 4x ACE-tRNA^Leu^ mNG plasmids (top) exhibit similar transfection efficiency to 16HBEge G542X-, R1162X- and W1282X-CFTR cells transfected with respective 4x ACE-tRNA mNG plasmids (bottom). Green traces represent mNG positive population of cells and gray traces represent cells transfected with empty plasmid without mNG expression cassette. (**B**) Representative FACS analysis histograms of non-trasfected (black trace) and empty plasmid transfected (grey) WT 16HBE14o- and 16HBEge G542X-, R1162X- and W1282X-CFTR cells. Overlap of non-transfected and empty plasmid transfected histograms indicate that transfection did not significant alter the gate used for FACS analysis (dashed line). (**C**) Real time qRT-PCR analysis of CFTR mRNA expression in WT 16HBE14o- (black checkered, *n* = 5), G542X-CFTR (orange, *n*= 2), R1162X-CFTR (red, *n* =2) and W1282X-CFTR cells (blue, *n* =2) transfected with empty mNG plasmid and sorted for positive mNG transduction. Data are presented as average ± SEM.

**Table S1. Percentage of mNG negative and positive cell populations for WT, G542X- R1162X- and W1282X-CFTR cells from FACS analysis.**

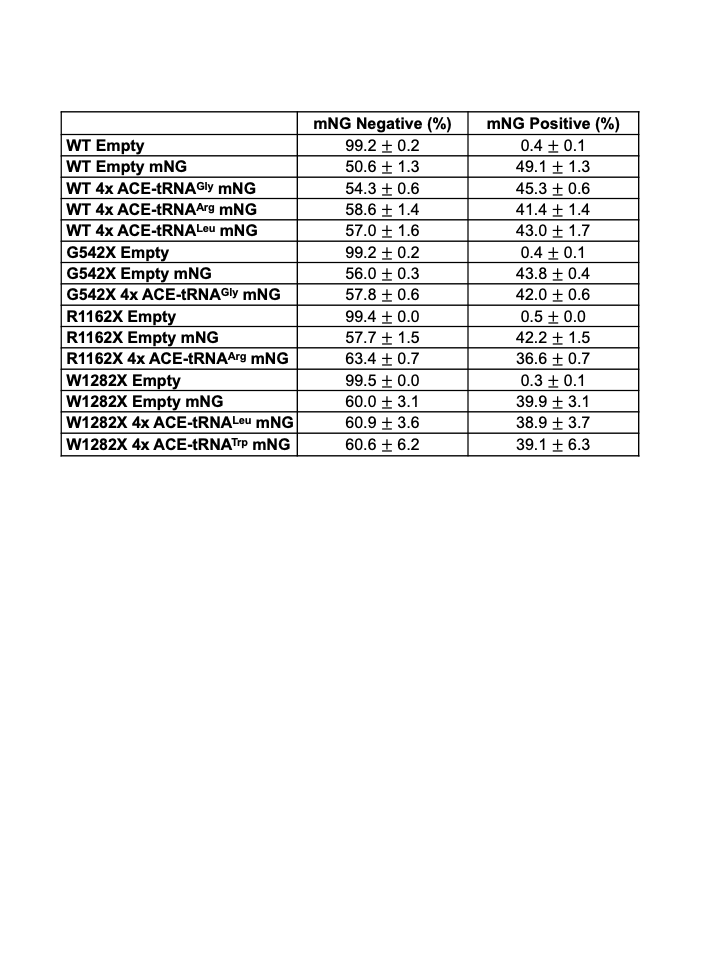

**Table S2. Sequences of oligonucleotides used in this study.**

| **Oligonucleotide Name** | **Oligonucleotide Sequence (5’-3’)** |
| --- | --- |
| EF1a pcDNA gibF | CGCGATGTACGGGCCAGATATACGCgctccggtgcccg |
| EF1a pcDNA gibR | AGCAGTGGGTTCTCTAGTTAGCCAGcctcacgacacctgaaatggaag |
| pcDNA prom gibR | GCGTATATCTGGCCCGTACATCG |
| pcDNA prom gibF | CTGGCTAACTAGAGAACCCACTGCTTAC |
| PROMgaF -> PmeI PB | GATCTGGAACGCTGTCTGAAGGATGCGATGTACGGGCCAG  ATATACGC |
| bGHgaR -> PmeI PB2 | TGACGGTATGGGCTCGCCTTAGTTTAAACAGCTGGTTCTTT  CCGCCTC |
| NLuc Bare/PEST -> EF1a gibF | CGGATCCACTAGTCCAGTGTGGTGGAATTCGCCACCATGGT  ATTCACACTCGAAG |
| NLuc Bare/Sec -> EF1a gibR | ATCAGCGGGTTTAAACGCTCTAGAGTCACTATTACGCCAGA  ATGCGTTCGCAC |
| NLuc PEST -> EF1a gibR | ATCAGCGGGTTTAAACGCTCTAGAGTCACTATTAGACGTTG  ATGCGAGCTGAAG |
| ArgTGAchr9.trna6/nointron F | CGTCGGCTCTGTGGCGCAATGGATAGCGCATTGGACTTCAA  ATTCAAAGGTTGTGGGTTCGAGTCCCACCAGAGTCG |
| ArgTGAchr9.trna6/nointron R | GGACCGACTCTGGTGGGACTCGAACCCACAACCTTTGAATT  TGAAGTCCAATGCGCTATCCATTGCGCCACAGAGCC |
| TrpTGAchr17.trna39 F | CGTCGGCCTCGTGGCGCAACGGTAGCGCGTCTGACTTCAGA  TCAGAAGGTTGCGTGTTCAAATCACGTCGGGGTCA |
| TrpTGAchr17.trna39 R | GGACTGACCCCGACGTGATTTGAACACGCAACCTTCTGATCT  GAAGTCAGACGCGCTACCGTTGCGCCACGAGGCC |
| GlyTGAchr19.trna2 F | CGTCGCGTTGGTGGTATAGTGGTTAGCATAGCTGCCTTCAAA  GCAGTTGACCCGGGTTCGATTCCCGGCCAACGCA |
| GlyTGAchr19.trna2 R | GGACTGCGTTGGCCGGGAATCGAACCCGGGTCAACTGCTTTG  AAGGCAGCTATGCTAACCACTATACCACCAACGC |
| GlnTAAchr17.trna14 F | CGTCGGTCCCATGGTGTAATGGTTAGCACTCTGGACTTTAAAT  CCAGCGATCCGAGTTCAAATCTCGGTGGGACCT |
| GlnTAAchr17.trna14 R | GGACAGGTCCCACCGAGATTTGAACTCGGATCGCTGGATTTAA  AGTCCAGAGTGCTAACCATTACACCATGGGACC |
| GluTAGchr13.trna2 F | CGTCTCCCACATGGTCTAGCGGTTAGGATTCCTGGTTCTAACCC  AGGCGGCCCGGGTTCGACTCCCGGTGTGGGAA |
| GluTAGchr13.trna2 R | GGACTTCCCACACCGGGAGTCGAACCCGGGCCGCCTGGGTTAG  AACCAGGAATCCTAACCGCTAGACCATGTGGGA |
| LeuTGAchr11.trna4 F | CGTCACCAGAATGGCCGAGTGGTTAAGGCGTTGGACTTCAGAT  CCAATGGATTCATATCCGCGTGGGTTCGAACCCCACTTCTGGTA |
| LeuTGAchr11.trna4 R | GGACTACCAGAAGTGGGGTTCGAACCCACGCGGATATGAATCC  ATTGGATCTGAAGTCCAACGCCTTAACCACTCGGCCATTCTGGT |
| EcoRVattR1/R2GAF | TACCGTCATTCATGCGGGATTATTCCAGTGTGGTGGAATTCTGC  AGATATCAAC |
| EcoRVattR1/R2GAR | TTACACTCGCTGACACTGATGTCGAGCGGCCGCCACTG |
| EF1a P3 ZraI gibF | CGAAAAGTGCCACCTGACCGATGTACGGGCCAGATATACGC |
| EF1a P3 ZraI gibR | AGTGCAGCATCATTGGGACAGCTGGTTCTTTCCGCCTC |
| CFTR probe | 56-FAM/TGCTTGTTT/ZEN/TAGAGTTCTTCTAATTATTTGGTATGTT  ACT/3IABkFQ |
| CFTR primer 1 | TTGCTGCTTGATGAACCCA |
| CFTR primer 2 | CATTGCTTCTATCCTGTGTTCAC |
| TBP probe | 5TET/TGATCTTTG/ZEN/CAGTGACCCAGCATCA/3IABkFQ |
| TBP primer 1 | GCTGTTTAACTTCGCTTCCG |
| TBP primer 2 | CAGCAACTTCCTCAATTCCTTG |
| puro probe | 56-FAM/TCGACATCG/ZEN/GCAAGGTGTGGGT/3IABkFQ |
| puro primer 1 | CCGATCTCGGCGAACAC |
| puro primer 2 | GTCACCGAGCTGCAAGAA |
| PB backbone probe | 56-FAM/CCAGATTGA/ZEN/GGGATACTTGCTGCCA/3IABkFQ |
| PB backbone primer 1 | GGCATGGGTACTTCCAAGAT |
| PB backbone primer 2 | CTCATAACCCAAGCCCTATCAG |
| RNase P probe | ABY/-TTCTGACCTGAAGGCTCTGCGCG/QSY |
| RNase P primer 1 | GAGCGGCTGTCTCCACAAGT |
| RNase P primer 2 | AGATTTGGACCTGCGAGCG |
